# Natural habitats are reservoirs of *Fusarium* lineages highly pathogenic on *Triticum aestivum*

**DOI:** 10.64898/2026.10.28.762564

**Authors:** Justine Mauriat, Benoit Lefebvre, Ludovic Bonhomme, Yves Martinez, Aurélie Le Ru, Fabienne Vailleau, Claudia Bartoli

## Abstract

Fungal plant diseases are a major cause of crop loss worldwide. Understanding the processes driving these diseases is therefore crucial to anticipate and control epidemics while improving crop yield production. Moreover, the new European « Green Deal » aims to reduce the use of pesticides, while, few alternatives are currently available on the market. There is therefore an urgent need to better track and anticipate fungal crop epidemics in the current context of fungicide reduction. In light of this, our study aims to understand *Fusarium* spill overs from non-agricultural habitats in order to better characterise potential non-host *Fusarium* reservoirs that may be implicated in the emergence of *Fusarium* diseases. To address this question, we isolated *Fusarium* lineages from natural populations of *Brassica rapa*. We characterised their phylogenomic affiliation and found that they display high diversity, with a population structure related to the locality of isolation. By inoculation assays, we identified *Fusarium* strains from non-agricultural habitats that were highly aggressive on wheat. In addition, we observed a strong effect of wheat variety on disease development. We monitored the *Fusarium* colonisation through a microscopy approach in both wheat and *Brassica* plants. We observed that while the fungus is mostly on root surface in *Brassica* plants, it spreads inter-cellularly in wheat roots with vascular colonisation in the wheat shoots. Finally, we showed that most aggressive *Fusarium* strains were isolated from Brittany, a French region where wheat is vastly cultivated.

## 1 Introduction

Plant disease epidemics have a major impact on society posing a significant threat to agricultural sustainability (Savary et al. 2019). Among plant diseases, fungal epidemics represent 30% of the disease outbreaks (Corredor-Moreno and Saunders 2020). Consequently, fungal plant pathogens are among the most destructive biotic constraints to agriculture, causing substantial reductions in crop yield and quality and resulting in worldwide economic losses estimated at hundreds of billions of US dollars annually (Dean et al. 2012; Fisher et al. 2012; Ojiambo et al. 2017).

Abiotic and biotic factors associated with conventional agricultural practices can enhance the dispersal, spread and emergence of plant pathogens. High vegetation density promotes pathogen transmission through direct contact between plants, particularly in mono-cultural cropping systems (Krnjaja et al. 2019). Fertilisation also contributes to disease development by increasing nutrient availability, which promotes plant growth, canopy density and leaf area, thereby creating favourable conditions for pathogen establishment and proliferation (Precigout et al. 2017; Krnjaja et al. 2019). In addition, intensive fertilisation can reduce plant association with beneficial microorganisms such as arbuscular mycorrhizal fungi, which not only enhance plant growth but also contribute to the stimulation of plant immunity processes (Oliveira Vieira et al. 2024). Finally, irrigation maintains high humidity and creates spatial-temporal bridges that increase pathogen survival, dispersal and sporulation in the case of phytopathogenic fungi (Anderson et al. 2004).

As for animal diseases, the emergence or re-emergence of plant diseases can result from the direct or indirect effects of multiple independent mechanisms. First, the intense use of fungicides in conventional agriculture is the major driver of disease emergence. Indeed, fungicides impose strong selective pressures on the pathogens thereby accelerating local adaptation and promoting the selection and spread of fungicide-resistant fungal lineages (Croll and McDonald 2017). Pathogens’ spill over from non-agricultural habitats to cultivated fields can also be mediated by human activities such as the expansion of cultivated areas (Srisakrapikoop et al. 2022). Also, the international movement of seeds between continents represents a global pathway for the introduction and spread of plant pathogens (Corredor-Moreno et al. 2019). Additionally, the water cycle can contribute to pathogen dispersal, as some fungi are capable of releasing and transporting their spores at long distances (Lagomarsino Oneto et al. 2020).

The selective processes acting on plant pathogens are not exclusively driven by human activities and environmental factors can exert strong selection on pathogen populations. For example, competition for nutrients and other resources, as well as exposure to antimicrobial compounds produced by resident soil microorganisms, can favour microbial traits that enhance pathogen survival. These ecological interactions may contribute to the local adaptation and persistence of pathogen populations within specific soil and habitat conditions (Weller et al. 2002). Mutations may also arise in fungal lineages that are already present in, or capable of colonizing crops, potentially leading to changes in genes associated with pathogenicity or aggressiveness. These newly acquired traits may subsequently be subjected to natural selection, depending on their effects on pathogen fitness in a given host or environment. In addition to mutation and recombination, horizontal gene transfer (Chellappan et al. 2026) and horizontal chromosome transfer (Ma et al. 2013), can contribute to the acquisition of pathogenicity and virulence-associated factors. For example, genetic regions carrying virulence-related genes can be transferred between *Fusarium* strains, potentially enabling previously non-pathogenic strains to infect new hosts or expanding the host range of existing pathogens (Friesen et al. 2006). These molecular mechanisms can then enhance the probability of pathogenic lineages to host jump and expand their host range thereby infecting new host genotypes or novel host species (Giraud et al. 2010; Kush et al. 2024).

Among the fungal phytopathogens, *Fusarium* regroups a large number of species able to infect a wide range of crops, as maize (*Zea mays*), potato (*Solanum tuberosum*) or wheat (*Triticum* spp.) (Armer et al. 2024). In wheat, *Fusarium graminearum*, *Fusarium culmorum,* and *Fusarium pseudograminearum* are known to be the causal agents of *Fusarium* crown rot (FCR) and the *Fusarium* head blight (FHB) diseases (Leplat et al. 2013; Scherm et al. 2013; Kazan and Gardiner 2018). *F. graminearum*, the principal causal agent of FHB, poses a serious threat to wheat and other small-grain cereals worldwide. This pathogen not only diminishes yield and grain quality but also contaminates harvests with harmful mycotoxins such as deoxynivalenol, representing significant threats to food safety and human and animal health (Xu et al. 2022). While most research on *F. graminearum* focuses on agricultural isolates, emerging evidence suggests that non-agricultural habitats, may harbour diverse potentially pathogenic *Fusarium* populations. For instance, *F. graminearum* has been consistently detected in wild grass hosts, with incidence correlated to rainfall and host density at the crop–wild interface (Fulcher et al. 2020). Moreover, *Fusarium* species, including *F. graminearum* and *F. culmorum,* isolated from asymptomatic weeds and non-graminaceous plants (*e.g. Brassica napus* and *Beta vulgaris*) have demonstrated pathogenicity toward wheat under greenhouse conditions (Zavtrikovienè et al. 2022). These findings point to a broader scenario in which non-agricultural reservoirs can serve as sources of inoculum for epidemics. Studies in other pathosystems reinforce this notion. For instance, research on *Pseudomonas syringae* has shown that environmental strains, isolated from snowpack and streams, harbour type III effector repertoires similar to those of tomato pathogens and are often equally aggressive (Monteil et al. 2016). Together, these observations underscore the imperative to broaden our surveillance and genomic analyses beyond agricultural fields.

To address the lack of information on the origin and evolution of pathogenic *Fusarium* lineages, this study characterises *Fusarium* strains isolated from non-agricultural habitats, both genetically and in terms of their pathogenicity. To this end, we isolated *Fusarium* lineages from natural *B. rapa* populations located in Italy and France and characterised their phylogenetic relationships and pathogenicity potential. We investigated whether *Fusarium* strains originated from non-agricultural habitats were able to infect bread wheat (*T. aestivum*). Specifically, we addressed the following hypotheses: (i) *Fusarium* lineages are host-specialised and this specialisation follows phylogenetic relationships; (ii) *Fusarium* lineages originating from non-agricultural habitats are less aggressive toward wheat than lineages isolated from wheat and they are able to infect a broad range of wheat varieties; and (iii) *Fusarium* lineages from non-agricultural habitats are less adapted and therefore less aggressive on ancient wheat varieties than modern varieties where they are more aggressive because of their maladaptation to recent host genotypes. In order to address these hypotheses, we characterised the phylogenetic affiliation of *Fusarium* strains isolated from *B. rapa,* evaluated their potential pathogenicity on wheat, using both ancient and modern (post World War II) wheat varieties. Finally, we investigated the cellular localization of these *Fusarium* isolates using microscopy, both on wheat and on their original *B. rapa* host.

## 2 Results

### Isolation and characterization of 102 *Fusarium* strains from non-agricultural habitats

Fungi were isolated from roots and rhizospheres of wild *B. rapa* populations collected in Sicily (Italy) and Brittany (France) between 2021 and 2022 (Maillet et al. 2025). These populations were distributed across highly diverse habitats that differed in their degree of anthropization and proximity to cultivated land. After four weeks of incubation of plated root and rhizosphere suspensions, 255 *Fusarium*-like isolates were recovered and purified onto Potato Dextrose Agar plates. *ITS1* sequencing of these 255 isolates revealed that only 148 belonged to the genus *Fusarium*. To further characterise the species identity of these strains, we additionally amplified the *RPB2* and *TEF-1*α loci and successfully sequenced both markers for 102 (52 and 50 isolates from France and Italy, respectively) out of the 148 strains (Table S1). A multilocus phylogenetic analysis based on *ITS1*, *RPB2*, and *TEF-1*α sequences was then performed (Data Set 1) using these 102 strains together with reference *Fusarium* genomes (Data Set 2) selected for their phylogenetic proximity to the 102 *Fusarium* strains. The resulting phylogenetic tree revealed a high diversity of the 102 *Fusarium* lineages isolated from non-agricultural habitats (Figure 1). Specifically, 30 strains belonged to the *F. culmorum/F. graminearum* species complex, which includes the *F. graminearum* strain MDC_Fg1 (Alouane et al. 2018) used in this study as reference strain for pathogenicity tests. Seventeen strains belonged to the *F. citri/F. equiseti/F. croceum* complex, 16 to the *F. acuminatum* cluster, and 12 to the *F. avenaceum* cluster. The remaining strains were less abundant and were distributed among the *F. flocciferum/F. iranicum*, *F. solani/F. stercicola*, *F. oxysporum/F. redolens*, and *F. sporotrichioides/F. brachygibbosum/F. cerealis/F. clavum* clusters (Figure 1).

**Figure 1.**
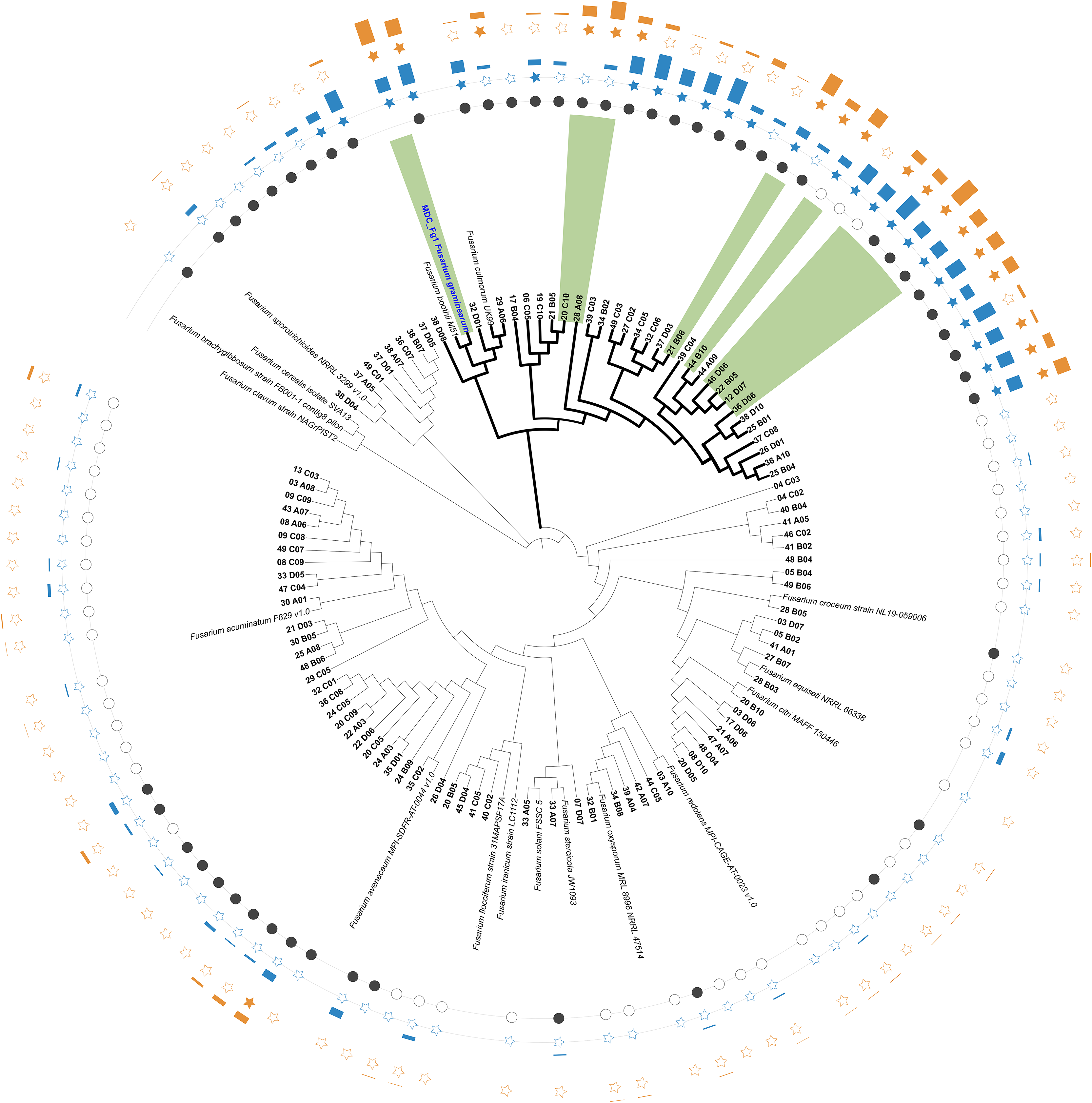
Phylogenetic tree of the 102 *Fusarium* strains isolated from rhizosphere and roots of *B. rapa* populations based on *ITS1*, *RPB2* and *TEF-1*α marker genes. Black dots correspond to *Fusarium* strains isolated in France, while white dots correspond to strains isolated in Italy. *F. graminearum*/*F. culmorum* complex is represented with bold branches and the reference MDC_Fg1 is visualized in bold blue. Blue histograms indicate the Least Squares means (LS-means) estimated on the level of pathogenicity of each strain on the Apache wheat variety in gnotobiotic conditions. Blue stars represent strains with a significant level of pathogenicity after an FDR test (*P* ≤ 0.05), white stars represent strains that not passed the FDR test. Orange histograms indicate the LS-means estimated on the level of pathogenicity of each strain on the Apache wheat variety under *in planta* conditions. Orange stars represent strains with a significant level of pathogenicity after FDR test (*P* ≤ 0.05). White stars represent strains that not passed the FDR test. *Fusarium* strains tested on the 20 wheat varieties are represented in green. The phylogentic tree was obtained with iTOL v7.6 (Letunic and Bork 2024)

The distribution of the 102 strains revealed a population structure associated with their habitat of origin (Figure 1). For instance, the *F. culmorum/F. graminearum* species complex comprised 26 strains isolated from French *B. rapa* populations, compared with three from Italian populations. The *F. sporotrichioides/F. brachygibbosum/F. cerealis/F. clavum* cluster was composed exclusively of strains isolated in France. Similarly, the *F. avenaceum* cluster included 11 strains from France and only one from Italy. In contrast, the *F. acuminatum* cluster consisted exclusively of strains isolated in Italy (Figure 1).

### Evaluation of the pathogenicity potential of the 102 *Fusarium* strains: linking the habitat of origin with strain aggressiveness on wheat

To evaluate their pathogenicity potential on wheat, the 102 *Fusarium* strains were inoculated on the two modern wheat varieties Apache and Paledor, in gnotobiotic and under *in planta* conditions (Data Sets 3,4,5). For each experimental condition, two temporal blocks were established, with four replicates per block. The reference *F. graminearum* MDC_Fg1 strain was included in each pathogenicity assay as a positive control. The disease scale for both gnotobiotic and *in planta* conditions is showed in Figure S1.

For the Apache variety, pathogenicity tests under gnotobiotic conditions showed that 57 of the 102 *Fusarium* strains tested were able to induce symptoms (Data Set 6). Additionally, the linear model showed a significant effect of the *Fusarium* strains’ factor when tested as fixed effect (*F* = 12.36, *P* ≤ 2.2e^10⁻¹^). However, only 24 *Fusarium* strains were considered significantly pathogenic with disease symptoms associated with FCR (*P* ≤ 0.05; Figures 1, 2A, Data Set 6). The 24 strains belonged to the *F. graminearum*/*F. culmorum* species complex, which includes the reference strain MDC_Fg1 (Figures 1, 2A, Data Set 6). Under *in planta* conditions, 51 out of the 102 strains were able to induce symptoms on the Apache wheat variety. We also observed a significant effect of the *Fusarium* strains’ factor considered as fixed effect (*F* = 12.66, *P* ≤ 2.2e^10⁻¹^). Specifically, 18 strains caused significant disease symptoms associated with FCR (*P* ≤ 0.05; Figures 1, 2B, Data Set 6). All pathogenic strains under i*n planta* conditions belonged to the F*. graminearum*/*F. culmorum* species complex, except for one strain phylogenetically related to the *F. avenaceum* cluster (Figure 1). Disease scores measured under *in planta* conditions were significantly correlated with that measured under gnotobiotic conditions (Spearman’s correlation, ρ = 0.42, *P* = 2.2e^10⁻¹^).

**Figure 2.**
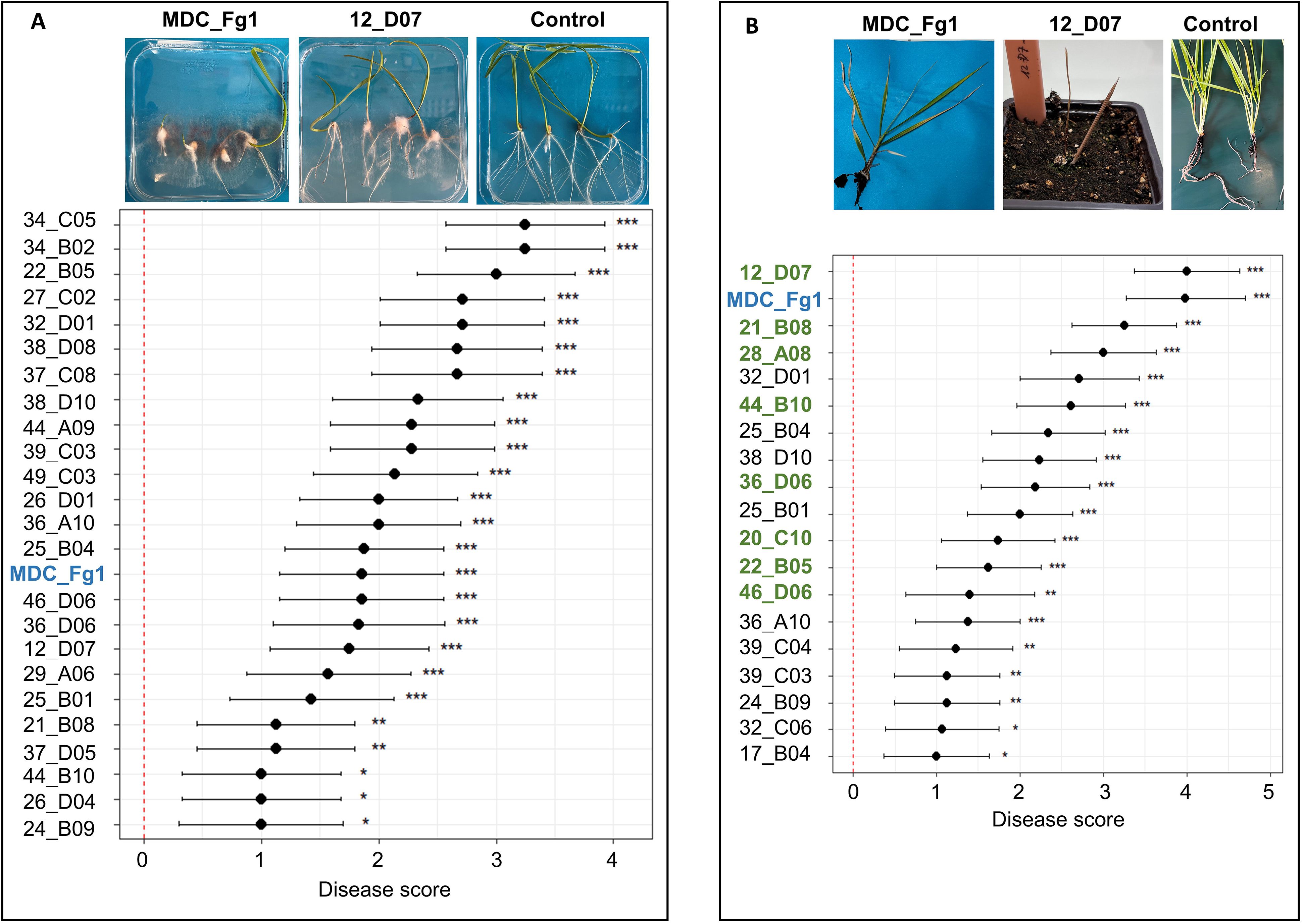
Pathogenicity tests of *Fusarium* strains inoculated on the Apache wheat variety. (A) LS-means estimated using disease score values (Figure S1) of the 25 *Fusarium* strains causing significant *Fusarium* crown rot (FCR) symptoms under gnotobiotic conditions. (B) LS-means estimated using disease score values (Figure S1) of the 18 *Fusarium* strains causing significant FCR symptoms under *in planta* conditions. Strains shown in bold green were selected for subsequent pathogenicity tests on 20 wheat varieties. (A-B) The MDC_Fg1 strain is indicated in bold blue. The red line indicates the negative control, corresponding to non-inoculated plants. Statistical significance is indicated as follows: *** *P* ≤ 0.0005, ** *P* ≤ 0.005, and * *P* ≤ 0.05. Representative disease symptoms caused by the MDC_Fg1 strain, the 12_D07 *Fusarium* strain, and the non-inoculated control are shown in the upper part, in the left, central, and right pictures, respectively.

On the Paledor wheat variety, 69 out of the 102 *Fusarium* strains were able to induce symptoms under gnotobiotic conditions, and the linear model showed a significant effect of the *Fusarium* strains’ factor when tested as fixed effect (*F* = 2.17, *P* ≤ 1.32e^10⁻07^). However, only 29 strains caused significantly FCR disease symptoms (*P* ≤ 0.05; Figure S2A, Data Set 6). Under *in planta* conditions, 11 *Fusarium* strains induced symptoms, of which only one caused significantly FCR disease symptoms (Figure S2B). As observed for the Apache variety, all pathogenic strains identified on Paledor originated from France, further supporting a potential geographical effect on the distribution of pathogenic *Fusarium* lineages. In addition, the pathogenic non-agricultural *Fusarium* strains were phylogenically located within the *F. graminearum*/*F. culmorum* species complex.

To further investigate the relationship between the habitat of origin and the pathogenicity of the *Fusarium* strains, we focused on the Apache variety, which exhibited a broader range of disease responses under *in planta* conditions. Using a linear model with the locality of isolation of the *Fusarium* strains as a fixed effect, we identified a highly significant effect of the country of origin on strain pathogenicity under gnotobiotic conditions (*F =* 125.87*, P* ≤ 2.2^e-16^; Figure 3A). In particular, Lande d’Apigné, Boël, and Bruz are three French localities, all located in the Brittany region, that were significantly associated with the pathogenicity *Fusarium* strains isolated from non-agricultural habitats (*F =* 9.75*, P* ≤ 2.2^e-16^ for the locality effect; Figure 3B). Among the Italian localities, Collesano was significantly associated with strain pathogenicity (Figure 3B). Under *in planta* conditions, the linear model also revealed a highly significant effect of the country of origin on strain pathogenicity (*F* = 70.39, *P* = 2.343^e-16^; Figure 3C). In particular, Lande d’Apigné, Boël, and Mordelles, the three French localities located in Brittany, were significantly associated with strain pathogenicity (*F* = 6.39, *P* ≤ 2.2^e-16^ for the locality effect; Figure 3D). Among the Italian localities, Gratteri and Collesano were associated with strain pathogenicity (Figure 3D). Altogether, these results indicate a strong effect of the strain origin on their pathogenicity potential on the Apache variety.

**Figure 3.**
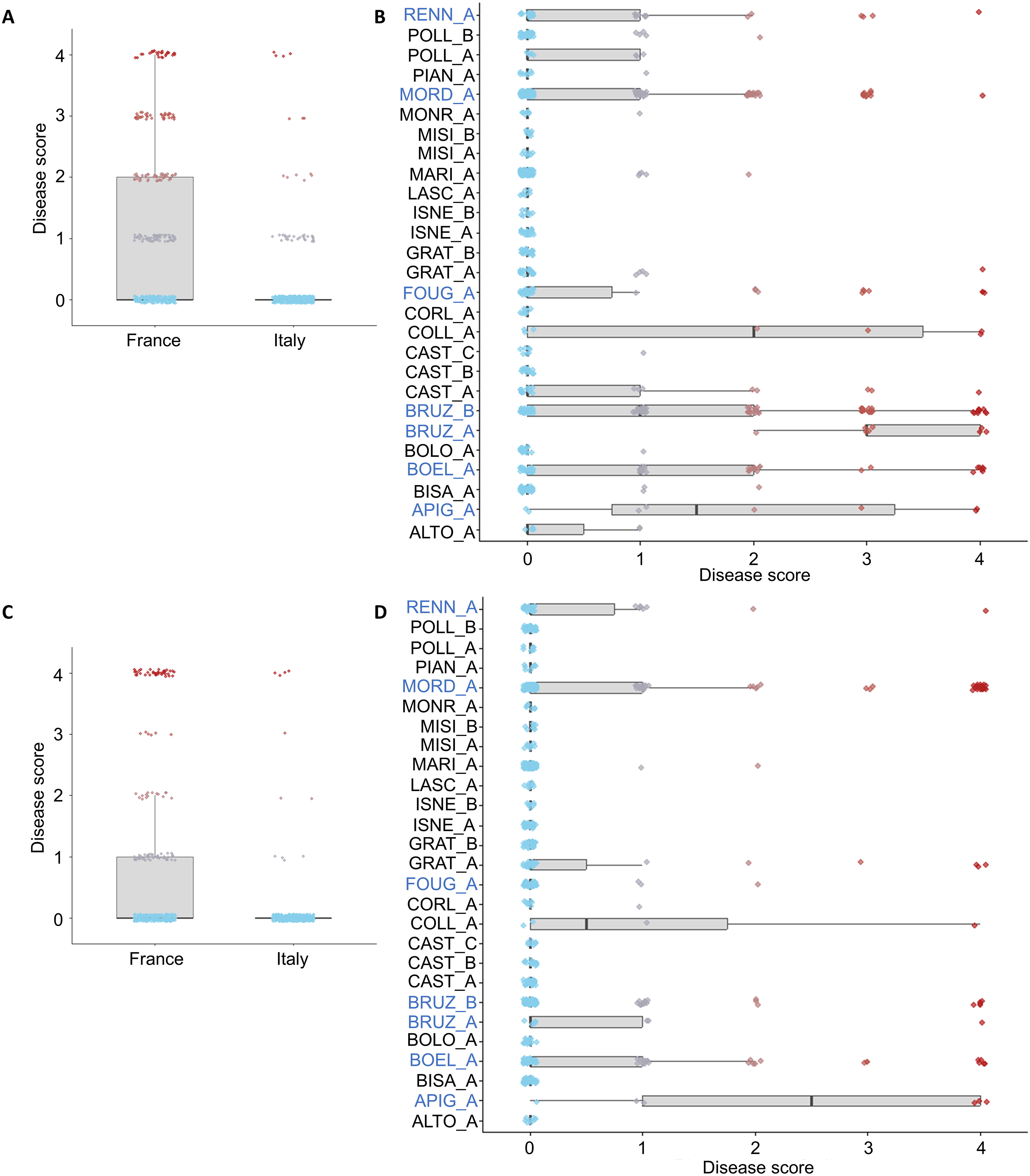
Effect of the country and locality of *Fusarium* strain origin on their aggressiveness on the Apache wheat variety. Disease scores by country of origin (A), and disease score by locality of origin (B) measured in gnotobiotic conditions. Disease score by country of origin (C), and disease score by locality of origin (D) measured under *in planta* conditions. French localities are indicated in blue. The disease score for each strain is represented by a dot and a colour gradient represent the disease score level (light blue = disease score 1 and dark red = disease score 4). Statistical significance is indicated as follows: *** *P* ≤ 0.0005, ** *P* ≤ 0.005, and * *P* ≤ 0.05

### Interaction between wheat varieties and *Fusarium* lineages on FCR disease

Based on pathogenicity assays conducted *in planta* on the Apache wheat variety, we selected eight pathogenic *Fusarium* strains originating from non-agricultural habitats and displaying a gradient of aggressiveness (Figure 2B). In particular, strain 12_D07 was selected as an aggressive strain, exhibiting a phenotype comparable to that of the reference MDC_Fg1. Strains 21_B08 and 28_A08 were selected as highly aggressive strains, 44_B10 and 36_D06 as moderately aggressive strains, and 20_C10, 22B_B05, and 46_D06 as weakly aggressive strains (Figure 2B). The eight *Fusarium* strains and the reference MDC_Fg1 were used to test the effect of wheat genetic variation on FCR disease development. For this, the wheat varieties were selected to cover both ancient and modern winter bread-wheat varieties. Eleven ancient varieties were selected to represent a gradient of genetic diversity and encompass 10 out of the 11 major global wheat genetic groups described by Balfourier et al. (2019). The nine modern varieties correspond to elite varieties cultivated in France over the last decade (Table S2).

To estimate the effect of wheat variety on wheat disease development, an initial analysis on the disease kinetic was performed. For this, the number of dead plants was recorded weekly over five weeks. Mortality data were recorded for each *Fusarium* strain and each wheat variety. We first analysed disease development by merging data for all plant varieties. We observed that the 12_D07 strain was responsible for the highest number of the plant deaths over time, with a total number of 128 dead plants out of 240 at the end of the experiment. This strain was followed by strain 22_B05, which caused 108 plant deaths out of 240, and the reference MDC_Fg1, which caused 89 deaths out of 240 (Figure 4A). The remaining *Fusarium* strains were associated with lower levels of plant mortality, ranging from 36 dead plants for strain 36_D06 and 15 dead plants for strain 20_C10 to (Figure 4A). We then analysed disease development across wheat varieties, which revealed a highly variable gradient of susceptibility among the 20 varieties (Figure 4B). No difference in mortality was observed between ancient and modern wheat varieties (Figure 4B). The ancient variety Manitoba showed the highest number of dead plants, with 37 out of 114 dead plants, followed by the modern variety Sy Mattis, with 35 deaths out of 114 plants (Figure 4B). In contrast, the ancient variety Cheyenne showed a comparatively higher level of resistance, with only 12 out of 114 plants dying during the experiment.

**Figure 4.**
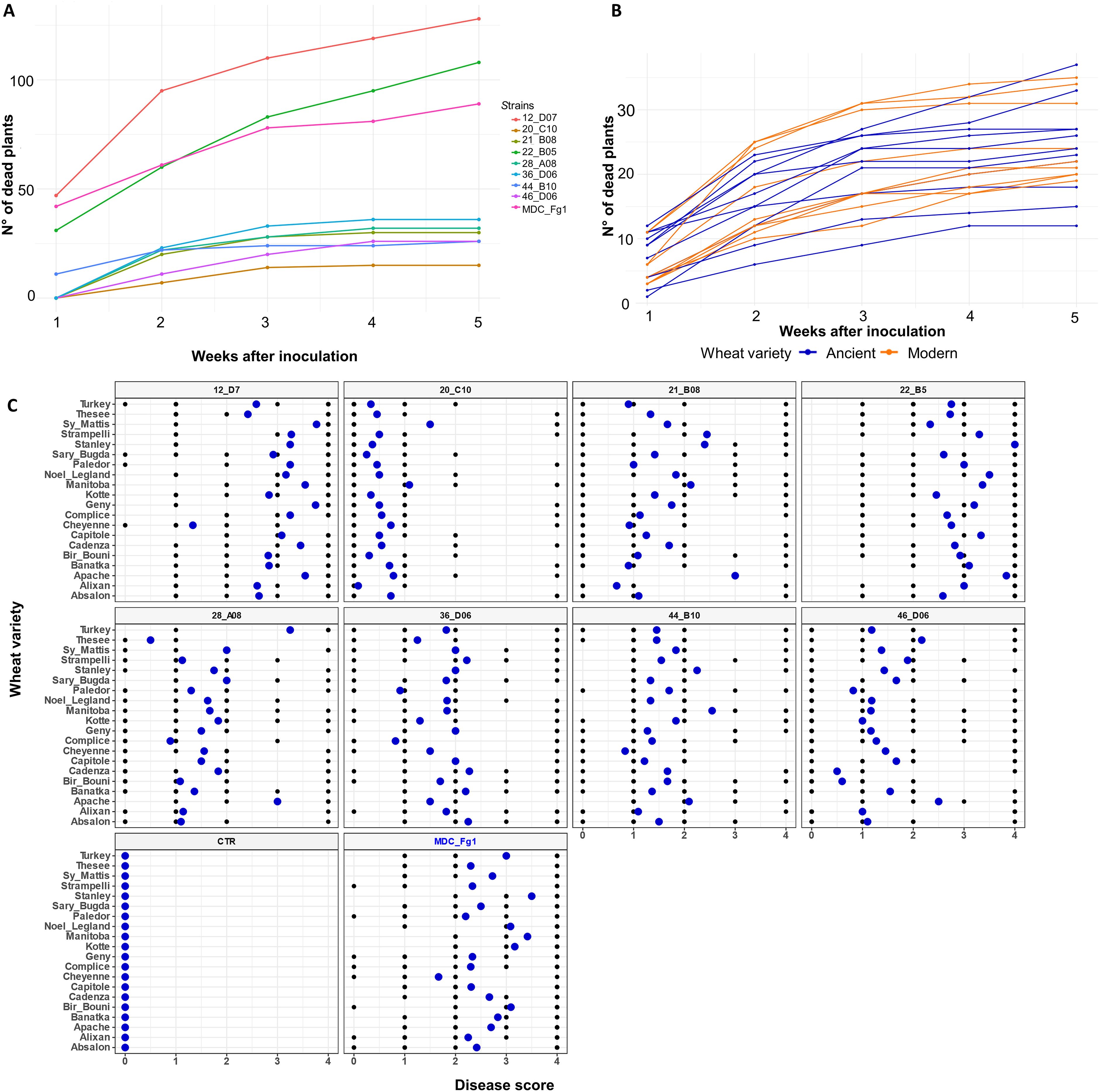
Evaluation of the effect of wheat genotypes on the development of *Fusarium* crown rot (FCR) caused by non-agricultural *Fusarium* strains and the reference MDC_Fg1. (A) Kinetics of FCR disease development, for each *Fusarium* strain on the merged 20 wheat varieties. (B) Kinetics of FCR disease development for each wheat variety for the nine merged strains. Blue lines represent ancient wheat varieties, whereas orange lines represent modern wheat varieties. (C) Distribution of FCR disease severity. Each panel represents the FCR disease severity associated with one *Fusarium* strain (*x*-axis) across the 20 wheat varieties (*y*-axis). Black points represent the distribution of the FCR disease for each wheat variety. Blue points represent the FDR corrected LS-means.

We then analysed the aggressiveness level of each strain and for each individual wheat variety considering FCR LS-means estimated using the disease scores at five weeks post-inoculation (Figure 4C). A linear mixed model, considering the *Fusarium* strain and the wheat variety as fixed effects, showed that the level of FCR disease was significantly influenced by the *Fusarium* strain factor (*F* = 118.74, *P* ≤ 2.2^e-16^, Figures 4C, S3A). When comparing with the negative control, the nine *Fusarium* strains induced significant FCR disease on all wheat varieties (Table S3, Figures S3A, S4A). On the other hand, all *Fusarium* strains from non-agricultural habitats differed significantly from one another (Table S4). In addition, pairwise comparisons within strains showed that the 12_D07 and 22_B05 strains were inducing a significantly higher level of disease compared to the other strains (Figures S3A, S4A, Table S5), with a level of aggressiveness similar to MDC_Fg1.

When considering the plant variety as fixed effect, the general linear mixed model showed a strong effect of the wheat variety on the development of the FCR disease (*F* = 4.09, *P* = 7.10^e-09^, Figures 4C, S3B, S4B). Specifically, the most susceptible wheat varieties were Apache, Manitoba and Stanley (*P* = 0.00035, *P* = 0.001 and *P* = 0.001 respectively, Figures 4C, S3B, S4B). It is worth to notice that Manitoba and Stanley are ancient varieties while Apache is a modern wheat variety. We also observed that the ancient variety Cheyenne displayed a resistant phenotype (*P* = 0.0084, Figure 4C, S3B, S4B).

To further assess the contribution of wheat variety in FCR establishment caused by *Fusarium* strains from non-agricultural habitats, we estimated the broad-sense heritability (H²) of disease response across the 20 wheat varieties. The heritability analysis showed that the response to FCR caused by the most aggressive non-agricultural strain, 12_D07, was largely explained by wheat variety, with an H² of 0.65, indicating that 65% of the phenotypic variation in disease response could be attributed to genetic differences among wheat varieties. For strains 28_A08, 44_B10, 21_B08, 36_D06, and the reference MDC_Fg1, H² values were 0.53, 0.51, 0.47, 0.39, and 0.20, respectively. In contrast, H² values were close to zero for strains 20_C10 and 46_D06. Altogether, these results indicate that wheat variety makes an important contribution to variation in FCR development caused by non-agricultural *Fusarium* strains, particularly for highly aggressive strains.

### Link between *Fusarium*-affiliated OTUs and *Fusarium* isolates in natural *B. rapa* populations

Given that several *Fusarium* strains isolated from non-agricultural habitats displayed high aggressiveness toward wheat, an important question is how prevalent these pathogenic lineages are in their natural habitats.

To address this question, we took advantage of previous metabarcoding data obtained from the same 26 *B. rapa* populations that were sampled for *Fusarium* isolation (Maillet et al. 2025). These data were used to identify OTUs taxonomically affiliated with *Fusarium* based on the *ITS1* marker. A total of six OTUs were identified as being affiliated with *Fusarium* species. Among them, OTU000003 was the most abundant, accounting for 2.3% of the total reads across the *B. rapa* samples and being detected in all *B. rapa* populations. OTU000016 accounted for 0.5% of the total reads and was detected in 18 out of the 26 *B. rapa* populations. In contrast, OTU000068 was detected exclusively in four French populations— Boël, Bruz, Landes d’Apigné, and Mordelles—with a relative abundance of 0.13% across all *B. rapa* samples. These four localities are all located in Brittany, where highly aggressive *Fusarium* lineages were isolated. The remaining OTUs, OTU00309, OTU00342, and OTU00848, were detected at very low relative abundances of 0.02%, 0.02%, and 0.007%, respectively.

To determine whether these *Fusarium* OTUs were phylogenetically related to the 102 *Fusarium* strains isolated by culturing, we performed a phylogenetic analysis based on *ITS1* (Figure 5). This analysis showed that the highly abundant OTU000003 was phylogenetically distant from the strains isolated, clustering with *F. clavum* (Figure 5). Moreover, none of the *Fusarium* strains isolated from clades closely related to *F. clavum* showed pathogenicity toward wheat (Figure 1). We also observed that OTU000016 and the three rare OTUs (OTU00309, OTU00342, and OTU00848), clustered within the *F. solani* clade (Figure 5). Several *Fusarium* strains isolated from *B. rapa* were closely related to this clade; however, none of these strains displayed pathogenicity toward wheat (Figure 1). In contrast, OTU000068, which was detected exclusively in the French populations, clustered within the *F. graminearum*/*F. culmorum* species complex (Figure 5). This clade includes all *Fusarium* strains identified as pathogenic toward wheat under both gnotobiotic and *in planta* conditions (Figure 1).

**Figure 5.**
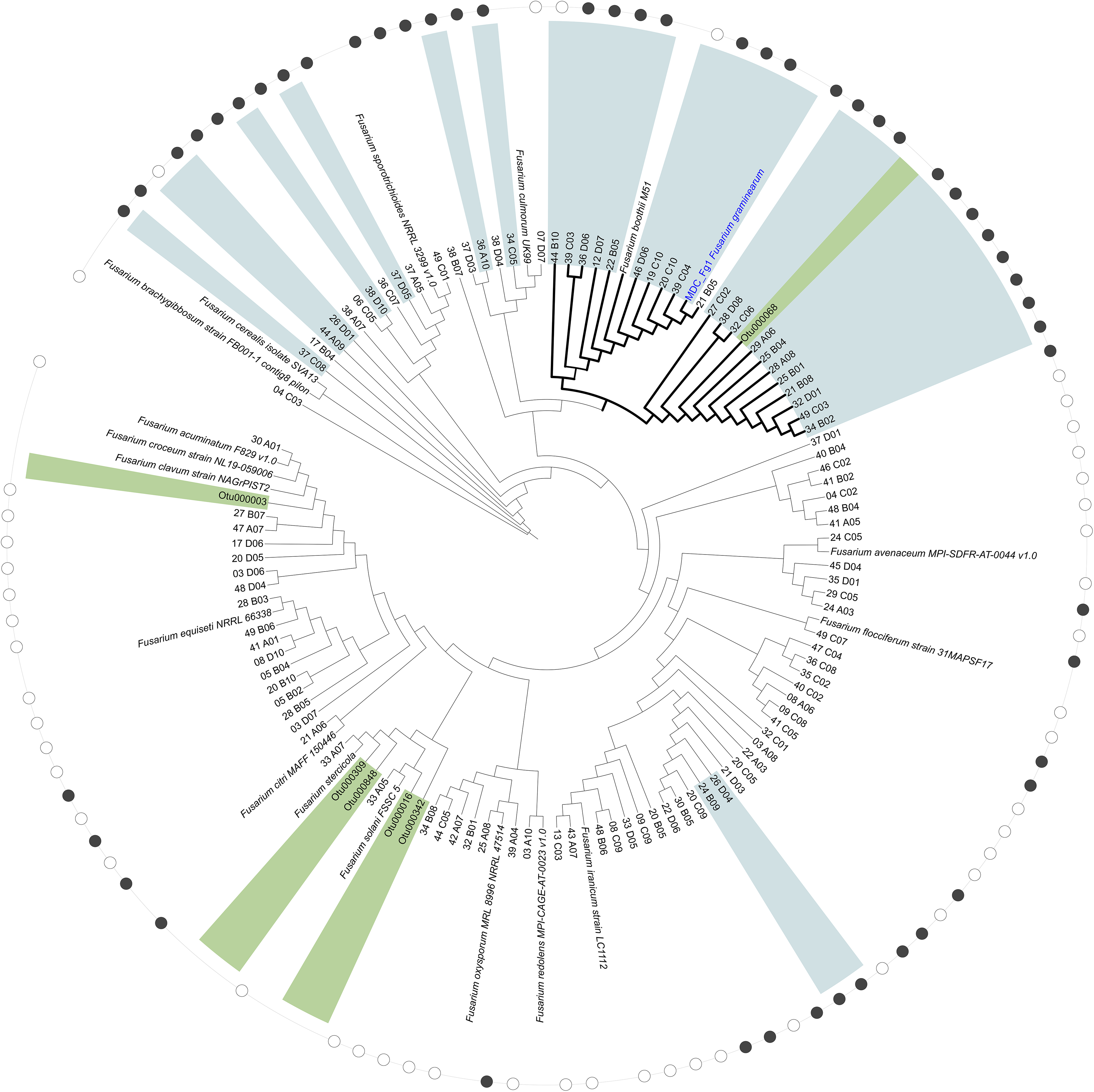
Phylogenetic tree based on an *ITS1* fragment comparing OTUs obtained by metabarcoding of natural *B. rapa* populations with *Fusarium* strains isolated from these populations. The six OTUs identified as taxonomically affiliated with *Fusarium* species are highlighted in green. *Fusarium* strains isolated from *B. rapa* populations and inducing FCR disease in wheat are highlighted in light blue. Black dots indicate *Fusarium* strains isolated from France, whereas white dots indicate strains isolated from Italy. Reference *Fusarium* species were included in the phylogenetic analysis.

Overall, these results suggest that wheat-pathogenic *Fusarium* strains belonging to the *F. graminearum*/*F. culmorum* species complex are present at low relative abundance in the natural *B. rapa* populations investigated. The closest metabarcoding OTU, OTU000068, accounted for only 0.13% of the total reads and was detected in only four French localities (Figures 5) consistent with the locality dependent aggressiveness of the isolated strains.

### *Fusarium in planta* localization in wheat and *B. rapa* plants

To investigate how the highly aggressive strain 12_D07 and the moderately aggressive strain 44_B10 colonize wheat tissues, we followed their *in planta* localization at 1, 2, and 3 weeks post-inoculation (wpi). For this, we used a fluorescent wheat germ agglutinin (WGA) staining approach combined with fluorescence microscopy. With this method, we observed *Fusarium* strains in green while the plant tissues are in blue due to autofluorescence (Figure 6). Both *Fusarium* strains are detected in the roots (Figures 6A-B) at 1, 2, and 3 wpi, with a predominant localization in intercellular spaces and to a lesser extent intracellularly (Figures 6D-E). In the crown, both strains were predominantly observed in the outer cell layers at 1, 2, and 3 wpi. In the stems, both strains were sporadically detected in the mesophyll and the vascular tissues at 2 and 3 wpi, with a more extensive colonization of the highly aggressive strain 12_D07 (Figure 6G) compared to the moderately aggressive strain 44_B10 (Figure 6F).

**Figure 6.**
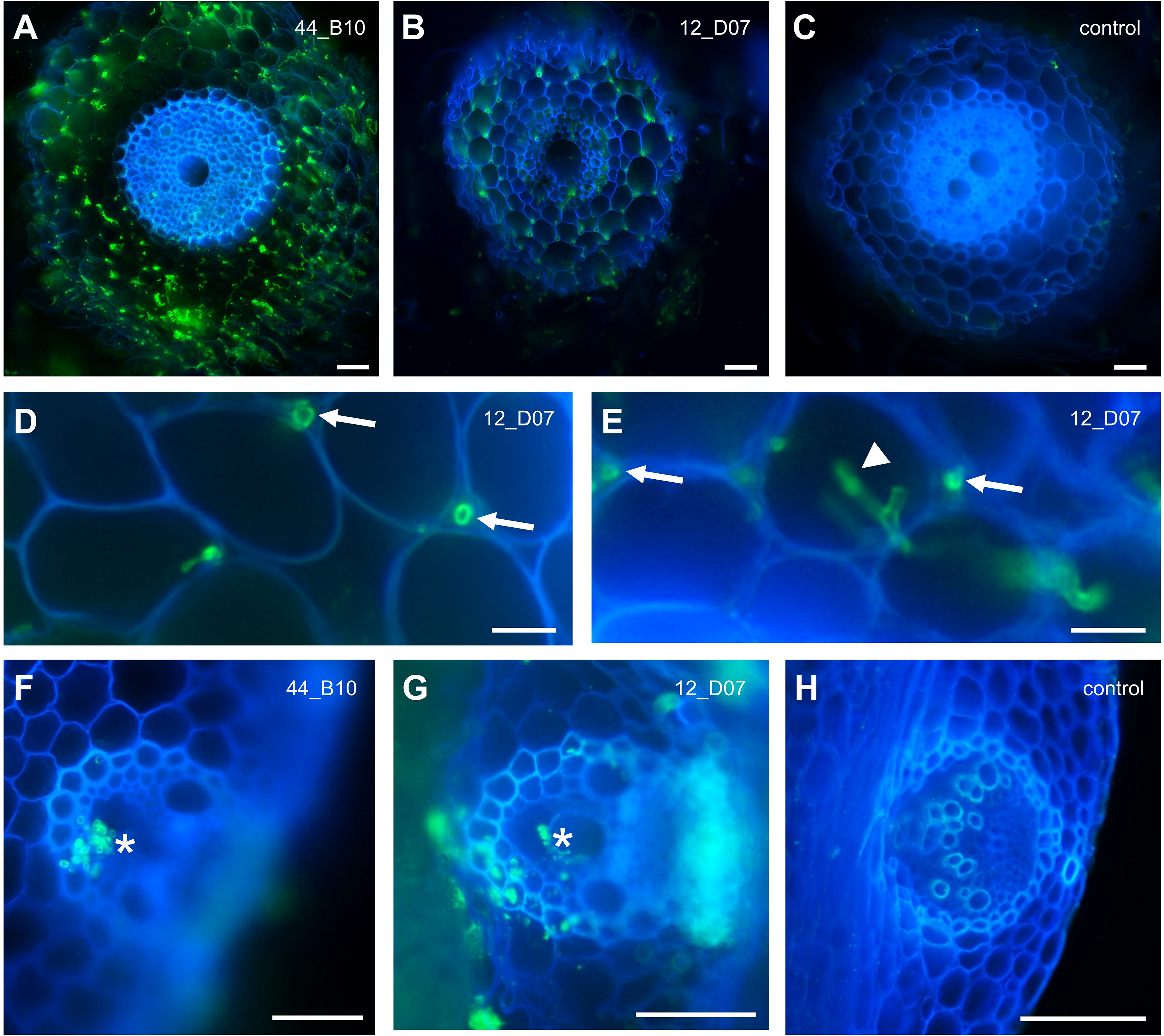
Fluorescence microscopy observations of *Fusarium* non-agricultural lineages after inoculation of the Manitoba wheat variety. Images of *Fusarium* colonization by the strains 44_B10 (A and F) and 12_D07 (B, D, E and G). Non-inoculated plants (C and H) were used as controls. The plant tissues are in blue (autofluorescence) and the *Fusarium* is visualized in green (wheat germ agglutinin staining). Both *Fusarium* strains were detected in roots (A-B; D-E) and in the stems (F-G). Arrows indicate intercellular colonisation; arrowheads indicate intracellular colonization and the stars indicate localization in the vascular tissues. Pictures were taken at 2 (A, C, D-F and H) and 3 (B and G) weeks post-inoculation. Bars represent 50 µm in A-C and F-H, and 25 µm in D-E.

To determine and characterise at which extent *Brassica* plants can host the two *Fusarium* strains, we used the same microscopic approach on the *B. rapa* BRUZ population, from which pathogenic French strains had been isolated, as well as the model fast-cycling *B. rapa* plants (Bartoli et al. 2026). We only observed a rare fungal root colonization of BRUZ population (Figure S5), and we could not detect a root colonization in the fast-cycling *B. rapa*. In both BRUZ and fast-cycling *B. rapa*, we observed few fungal structures into the stem, predominantly at the tissue surface.

## 3 Discussion

Fungal pathogens represent a major threat to agricultural production, with direct and indirect consequences for human health through yield losses and mycotoxin contamination (Dean et al. 2012; Fisher et al. 2012). Non-agricultural habitats are increasingly recognized as important reservoirs of potentially fungal phytopathogens (Bardin et al. 2018). Previous studies have shown that pathogenic strains originating from non-agricultural reservoirs can exhibit broader host ranges than strains associated with agricultural systems (Bartoli et al. 2015; Morris et al. 2008). Conversely, crop-associated strains may become increasingly specialized through co-evolution with their hosts, sometimes losing effectors and pathogenicity determinants involved in the colonization of a broad range of plant species (Monteil et al. 2016).

In the context of phytopathogens emergence, our initial hypothesis was that *Fusarium* lineages isolated from natural populations of *B. rapa* would be able to infect a broad range of wheat varieties, through the action of generalist pathogenicity determinants, because they had not co-evolved with particular wheat varieties. Also, our hypothesis was that these strains would display lower aggressiveness than epidemic strains originating from agricultural environments. However, our results only partially support this hypothesis. Several *Fusarium* strains isolated from natural *B. rapa* populations were pathogenic toward a broad panel of wheat varieties, including both modern and ancient varieties. Moreover, several strains displayed levels of aggressiveness exceeding those of the well characterized *F. graminearum* MDC_Fg1 reference strain, which was originally isolated from wheat and is known to cause severe FCR and FHB disease (Alouane et al. 2018). These results contrast with previous studies of pathogens isolated from non-agricultural environments, which have generally reported having a reduced pathogenicity associated with differences in effector repertoires and adaptation to alternative hosts (Morris et al. 2008; Monteil et al. 2013). Our results therefore indicate that non-agricultural origin does not necessarily imply reduced pathogenicity toward an agricultural host. Importantly, all strains that were significantly pathogenic toward wheat belonged to the *F. graminearum*/*F. culmorum* species complex. This strong phylogenetic association suggests that the ability to cause FCR on wheat appears to be a specialisation within particular evolved *Fusarium* lineages, rather than a random distribution across *Fusarium* lineages hosted in natural *B. rapa* populations. This result is consistent with the hypothesis that members of the *F. graminearum*/*F. culmorum* complex are the major causal agents of FCR in wheat and share pathogenicity determinants that facilitate infection of their host (Dyer et al. 2009; Winter et al. 2018). Genome studies of *F. graminearum* have identified numerous genes associated with host infection, including secreted proteins and other candidate effectors that may contribute to host colonization and FHB disease establishment in wheat (Brown et al. 2012; Rocher et al. 2022; Ayilalath et al. 2026). Studies in *Fusarium oxysporum* have demonstrated that lineage-specific pathogenicity chromosomes and associated virulence factors can strongly influence host specificity and disease development (Ma et al. 2013). Our phylogenetic and pathogenicity data therefore support an association between evolutionary lineage and pathogenic potential, although the presence of specific effectors or other pathogenicity determinants remains to be demonstrated experimentally in our *Fusarium* strains.

The geographical distribution of the pathogenic *Fusarium* strains identified in this study let us hypothesise that agricultural and non-agricultural habitats may be connected, as demonstrated in the case of zoonoses outbreaks (Perfecto et al. 2023). Indeed, most of the highly aggressive *Fusarium* strains were isolated in Brittany, a region characterised by intensive agriculture and cereal production, including wheat (Iachkine et al. 2026). This geographical enrichment raises the possibility that some of the *Fusarium* isolates may have an agricultural origin and subsequently colonised wild plant populations. Such a scenario would be consistent with the hypothesis that some pathogens detected in natural habitats represent “escaped” lineages originating from agroecosystems that subsequently established associations with wild plant populations (Power and Mitchell 2004). Agricultural landscapes are highly connected to semi-natural habitats, facilitating microorganism movement between crops and non-cultivated vegetation through wind, crop residues, insects, animals, and human activities (Berg et al. 2017). The capacity of *Fusarium* spores to disperse aerially over long distances could further facilitate movement between agricultural fields and surrounding habitats (Hoffmann et al. 2021). The comparison between France and Italy provides further support for a possible influence of agricultural context in the spread of pathogenic *Fusarium* lineages outside agricultural fields. The pathogenic strains originating from Sicily, where wheat cultivation is absent from the sampled environments, were less aggressive compared to MDC_Fg1 and to strains isolated in Brittany. This geographical pattern is compatible with the results reported for *P. syringae*, for which strains originating from natural habitats that were not associated with crop escape pathogen lineages, were generally less aggressive than epidemic crop-associated strains (Demba Diallo et al. 2012). However, fungal dispersal (Nabeela and Uppuluri 2025), environmental selection (Farrer et al. 2025), gene flow (Ropars et al. 2018), and differences in host related plant communities (Liu et al. 2021) could all contribute to the observed geographical structure. Nevertheless, the coexistence of highly aggressive and less aggressive *Fusarium* lineages within natural *B. rapa* populations suggests that non-agricultural habitats can harbour substantial variation in pathogenic potential.

Previous metabarcoding results provided an additional perspective on the role of non-agricultural habitats on the spread of fungal pathogenic lineages. Indeed, the OTU000068 phylogenetically associated with the *F. graminearum*/*F. culmorum* complex and corresponding to the lineage containing the wheat-*Fusarium* pathogenic strains was detected at a lower relative abundance, but was restricted to four Brittany populations. Together, these results suggest that the natural *B. rapa* populations surrounding agricultural landscapes may harbour members of the same phylogenetic lineage as wheat-pathogenic *Fusarium*. However, because OTU abundance represents relative sequencing read abundance rather than absolute fungal biomass (Tkacz et al. 2018), the low abundance of OTU000068 should not be interpreted as evidence of a low biological abundance of the corresponding fungus. Rather, its restricted distribution and phylogenetic association with pathogenic isolates provide evidence for a geographically structured reservoir of *Fusarium* lineages with potential relevance to wheat disease.

The ability of some non-agricultural *Fusarium* strains to cause severe disease and colonize wheat was further supported by microscopy observations. Two selected pathogenic stains with different levels of aggressiveness were able to penetrate wheat roots and colonize internal tissues, demonstrating that the infection was not restricted to the initial seed inoculation site, with a progressive colonization of aerial tissues. This pattern is consistent with previous works showing that *F. graminearum* can systemically colonize wheat (Jansen et al. 2005; Beyer et al. 2014). These observations constitute additional elements demonstrating that several strains isolated from natural *B. rapa* populations are *bona fide* wheat pathogens, colonizing the whole plant, reaching tissues far from the initial root inoculation site. This is consistent with previous work where *F. graminearum* was shown to colonize wheat ears through intercellular and intracellular growth, with extensive tissue colonization during advanced infection (Brown et al. 2010).

The variability observed among wheat varieties indicates that host genetic diversity remains a major determinant of susceptibility/resistance, even when plants are challenged with highly aggressive strains originating from non-agricultural habitats. Our work demonstrates that maintaining broad genetic diversity in breeding programs is crucial, and may be more informative than relying on a simple ancient *vs* modern distinction. Finally, our findings have implications for disease surveillance and sustainable disease management. Indeed, surveillance strategies focused exclusively on agricultural fields may overlook pathogenic lineages maintained in surrounding vegetation or semi-natural habitats.

## 4 Experimental procedures

### 4.1 Isolation and characterization of *Fusarium* strains from natural populations of *B. rapa* and fungal culture conditions

To isolate *Fusarium* lineages from 26 natural *B. rapa* populations collected in Italy (Sicily) and France (Brittany and Pays de la Loire) (Maillet et al. 2025), three plants per population were collected from the field. Roots and rhizospheres were separated *in situ* and placed in sterile 5-mL tubes. Samples were then transferred to the laboratory and stored in 40% glycerol at −20°C until processing. For fungal isolation, lateral roots were macerated in 1 mL of sterile distilled water using a scalpel. One gram of rhizosphere soil was also suspended in 1 mL of sterile distilled water and thoroughly mixed. Root macerates and rhizosphere suspensions were then plated onto several culture media (El-Oraiby, unpublished data), all supplemented with antibiotics to inhibit bacterial growth: 5 mg/L gentamicin, 300 mg/L streptomycin, 100 mg/L chloramphenicol, and 100 mg/L penicillin. Plates were incubated for 4–6 weeks at 15°C in the dark.

*Fusarium*-like mycelia were subsequently transferred to Potato Dextrose Agar (PDA; Merck, 1.01030) and sub-cultured until pure isolates were obtained. All *Fusarium*-like strains were then maintained in 5-mL slanted PDA tubes at 4°C. A strain of *F. graminearum*, called MDC_Fg1, isolated from symptomatic wheat plants and previously shown to cause both FCR and FHB in wheat (Alouane et al. 2018), was included as a positive control in the pathogenicity assays. The list of all strains isolated from *B. rapa* populations used in the study is available in Table S1. For all experiments described in this study, *Fusarium* strains were retrieved from the working collection, cultured on PDA Petri dishes, and incubated at 20°C in the dark for one week to obtain fresh and homogeneous mycelial material for each plant experiment. The whole experimental design is illustrated in Figure S6. *Fusarium* strains, including the reference MDC_Fg1, were characterized by sequencing three molecular markers: the translation elongation factor 1-α (*TEF-1*α), internal transcribed spacer (*ITS1*), and RNA polymerase II subunit (*RPB2*) genes (Table S6; Supplementary Text). Detailed experimental procedures for molecular characterization and phylogenetic analyses are provided in the Supplementary Text.

### 4.2 Plant material

We used a core collection of 20 bread wheat (*T. aestivum*) varieties (Table S2). Among these varieties, nine are modern varieties (elite varieties bred post-World War II) that were cultivated in France over the last decades, whereas 11 are ancient varieties. The 20 wheat varieties were produced at the experimental unit (UE APC) of INRAE in Castanet-Tolosan. All *in planta* experiments were performed in a growth chamber under controlled conditions (25°C day/22°C night, 16-h photoperiod, and 55% relative humidity) using non-sterile fertile potting soil (Proveen Rempotage 2). To avoid the presence of *Fusarium* strains from our stock of seeds, we performed seed sterilization before each experimentation in accordance with the protocol described in Bartoli et al. (2022). Then, seeds were incubated at 4°C during 2 or 3 days for the breaking of dormancy, then at 25°C for germination.

### 4.3 Inoculation of *Fusarium* strains under gnotobiotic and *in planta* conditions and evaluation of FCR disease score

For experiments under gnotobiotic conditions, seeds of the Apache and Paledor wheat varieties were surface-sterilized Bartoli et al. (2022). Four pre-germinated 3-day old seeds were then placed on sterile square Petri dishes (12 × 12 cm) containing M medium (Bartoli et al. 2022). A fungal suspension was prepared for each of the 103 *Fusarium* strains (102 non-agricultural strains and the reference MDC_Fg1). For each strain, 2 mL of sterile distilled water was added to a fungal culture grown on a 5.5-cm-diameter Petri dish. The fungal mycelium was suspended by gently scraping the culture surface with a sterile pipette tip, and 20 µL of the resulting suspension was deposited close to the wheat roots. To protect the roots from the light, the bottom of each Petri dish was wrapped in aluminium foil. FCR symptoms were assessed one week after inoculation using a disease scale ranging from 0 (no symptoms) to 4 (plant death) (Figures S1, S6). Two independent experiments, corresponding to two temporal blocks, were conducted, with four plants per block.

For experiments under *in planta* conditions. Apache and Paledor seeds were surface-sterilized as described above, and four pre-germinated seeds were transferred in individual pots containing non-sterile soil. A 1 × 1 cm plug of fresh fungal mycelium was then placed next to each wheat root using a sterile toothpick. Four weeks after inoculation, disease symptoms were assessed for each plant using a 0–4 disease scoring scale (Figures S1, S6).

To follow FCR development on 20 wheat varieties, eight *Fusarium* strains and the reference MDC_Fg1 were inoculated onto 12 plants of each of 20 wheat varieties (Table S2), resulting in a total of 240 plants tested for each fungal strain. The experiment was conducted using a split-plot design to limit random effects, and to assess the effects of wheat variety (fixed factor) and fungal strain (fixed factor) on the development of FCR. For each wheat variety, six non-inoculated plants were added as negative controls. More specifically, for each fungal strain, nine spatial blocks were established, with one or two plants of each wheat variety randomly distributed within each block. The blocks were randomized every three days throughout the five-week experiment. A nutrient solution (modified Hoagland solution; Bartoli et al. 2026) was supplied every two weeks. Two methods were used to assess disease development in this experiment: (i) the number of dead plants was recorded weekly for each fungal strain and wheat variety to monitor disease progression over time; and (ii) disease scores were measured based on FCR symptoms five weeks after inoculation with a disease index scale ranging from 0 (healthy plants) to 4 (dead plants) (Figure S1).

### 4.4 Statistical analysis and *Fusarium* localization in wheat tissues

Statistical analysis and microscopy procedures to evaluate the colonization level of the two non-agricultural strains 12_D07 and 44_B10 in wheat and *B. rapa* tissues are detailed in Supplementary Text.

## Supporting information

Data Sets

## 5 Acknowledgements

We particular thank Kehaulani An Tai and Chrystel Gibelin for their kind help in plant experiments. We acknowledge the TRI-FRAIB imaging facility, member of the national infrastructure France-BioImaging supported by the French National Research Agency (ANR-24-INBS-0005 FBI BIOGEN). We thank Gilles Tison and the “*Unité Expérimentale d’AgroEcologie et Phénotypage des Cultures*” of INRAE Occitanie-Toulouse for seed multiplication.

## CRediT author contributions

**Justine Mauriat**: methodology, data curation, formal analysis, writing original draft, review and editing. **Benoit Lefebvre**: methodology, review and editing. **Ludovic Bonhomme**: review and editing. **Fabienne Vailleau**: writing – original draft, conceptualization, supervision, review and editing. **Yves Martinez**: methodology. **Aurélie Le Ru**: methodology. **Claudia Bartoli**: data curation, supervision, project administration, resources, investigation, funding acquisition, conceptualization, formal analysis, validation. writing – review and editing.

## Supporting Information legends

Supporting Information are all available in the supporting file including:

## Supplementary Figures

**Figure S1. Disease scale used in gnotobiotic and *in planta* conditions to evaluate the aggressiveness of *Fusarium* strains**. Plants were recorded one week post inoculation in gnotobiotic conditions (A), while disease records were measured 4 weeks after inoculation under *in planta* conditions (B).

**Figure S2. Pathogenicity tests of *Fusarium* strains inoculated on the Paledor wheat variety.** (A) LS-means estimated using the disease score values of the 30 *Fusarium* strains causing significant Fusarium crown rot (FCR) symptoms under gnotobiotic conditions. The reference MDC_Fg1 is indicated in bold blue. The vertical red line indicates the negative control, corresponding to plants not inoculated with *Fusarium*. (B) LS-means for the *in planta* conditions pathogenicity test. For both tests only strains showing visible FCR symptoms are showed. Statistical significance is indicated as follows: *** *P* ≤ 0.0005, ** *P* ≤ 0.005, and * *P* ≤ 0.05.

**Figure S3. Pathogenicity tests of eight *Fusarium* strains from non-agricultural habitats and the reference MDC_Fg1 on 20 wheat varieties.** (A) Box plot of the level of FCR disease (*y*-axis) distributed over the nine strains tested and the non-inoculated control (CTR) (*x*-axis). (B) Box plot of the level of FCR disease (*y*-axis) distributed over the 20 wheat varieties (*x*-axis).

**Figure S4. Plots of the LS-means estimated on the disease score values of eight *Fusarium* strains from non-agricultural habitats and the reference MDC_Fg1 on 20 wheat varieties.** (A) LS-means plot of the level of FCR disease (*y*-axis) distributed over the nine strains tested and the non-inoculated control (CTR) (*x*-axis). (B) LS-means plot of the level of FCR disease (*y*-axis) distributed over the 20 wheat varieties.

**Figure S5. Microscopy observation of *Fusarium* non-agricultural lineages in the *B. rapa* origin host.** Images of the colonization by the *Fusarium* strain 44_B10 (A-B) at 1-week post-inoculation (wpi) on *B. rapa*. After fungal staining with WGA-FITC, root sections were performed using a microtome and imaged by using a fluorescence microscope with a DAPI (blue, plant autofluorescence) and a GFP filter (green, fungal staining). Non-inoculated plants (C) were used as controls. Bars represent 50 µm.

**Figure S6. Illustration of the experimental design adopted in our work**. *Fusarium* isolated strains were phylogenetically characterized and their pathogenicity was evaluated in gnotobiotic and planta conditions. The 103 *Fusarium* strains referred to the 102 non-agricultural strains and the reference MDC_Fg1. A set of 8 non-agricultural strains and the reference *F. graminearum* MDC_Fg1 strain were evaluated for their pathogenicity on a wheat core collection of 20 varieties.

## Supplementary Tables

**Tables S1**. List of *Fusarium* strains isolated from *Brassica rapa* natural populations used in this study.

**Table S2**. List of 20 wheat varieties used to test the effect of host genotype of Fusarium crown rot (FCR) disease of eight non-agricultural *Fusarium* strains and the reference *F. graminearum* MDC_Fg1. NA: not available.

**Table S3**. Pairwise comparisons between the nine *Fusarium* strains and the negative control (CTR) on 20 wheat varieties five weeks post-inoculation.

**Table S4**. Pairwise comparisons between the nine *Fusarium* strains and the reference MDC_Fg1 used as a positive control on 20 wheat five weeks post-inoculation.

**Table S5**. Pairwise comparisons between *Fusarium* strains during the experimental assays testing the development of FCR disease in the 20 wheat varieties, at five weeks post-inoculation.

**Table S6**. Primers used to amplify housekeeping genes in *Fusarium* strains.

## Supplementary Text

Supplementary experimental procedures

Supplementary references

## Data availability statement

Data to support our findings are available in the Data Set file including:

**Data Set S1.** Sequences of the *TEF-1*α, *RPB2* and *ITS1* obtained on the 102 *Fusarium* non-agricultural isolates and the reference MDC_Fg1 *F. graminearum* strain.

**Data Set S2**. Sequences of the *TEF-1*α, *RPB2* and *ITS1* from the reference *Fusarium* species obtained in the GenBank and used to affiliate *Fusarium* non-agricultural isolates.

**Data Set S3**. Raw data of pathogenicity values after the inoculation on the Apache wheat variety of the 102 *Fusarium* and the reference MDC_Fg1 in both gnotobiotic and *in planta* conditions.

**Data Set S5**. Raw data of pathogenicity values after the inoculation on the Paledor wheat variety of the 102 *Fusarium* and the reference MDC_Fg1 under *in planta* conditions.

**Data Set S5**. Raw data of pathogenicity values after the inoculation on the Paledor wheat variety of the 102 *Fusarium* and the reference MDC_Fg1 in gnotobiotic conditions.

**Data Set S6.** Statistical outputs obtained on the pathogenicity assays after inoculation of the 102 *Fusarium* non-agricultural strains in gnotobiotic and under *in planta* conditions.

**Data Set S7**. Raw data of the disease scores of the 20 wheat varieties measured 5 weeks after inoculation with the eight *Fusarium* strains.

**Data Set S8**. Raw data of the number of dead plants per *Fusarium* strain measured along 5 weeks after inoculation.

**Data Set S9**. Raw data of the number of dead plants of the 20 wheat varieties per wheat variety along 5 weeks after inoculation.

