## Supplementary Information for "Natural habitats are reservoirs of *Fusarium* lineages highly pathogenic on *Triticum aestivum*"

**This supplementary file includes**:

Supplementary Figures

Supplementary Tables

Supplementary Text

**Supplementary Figures**

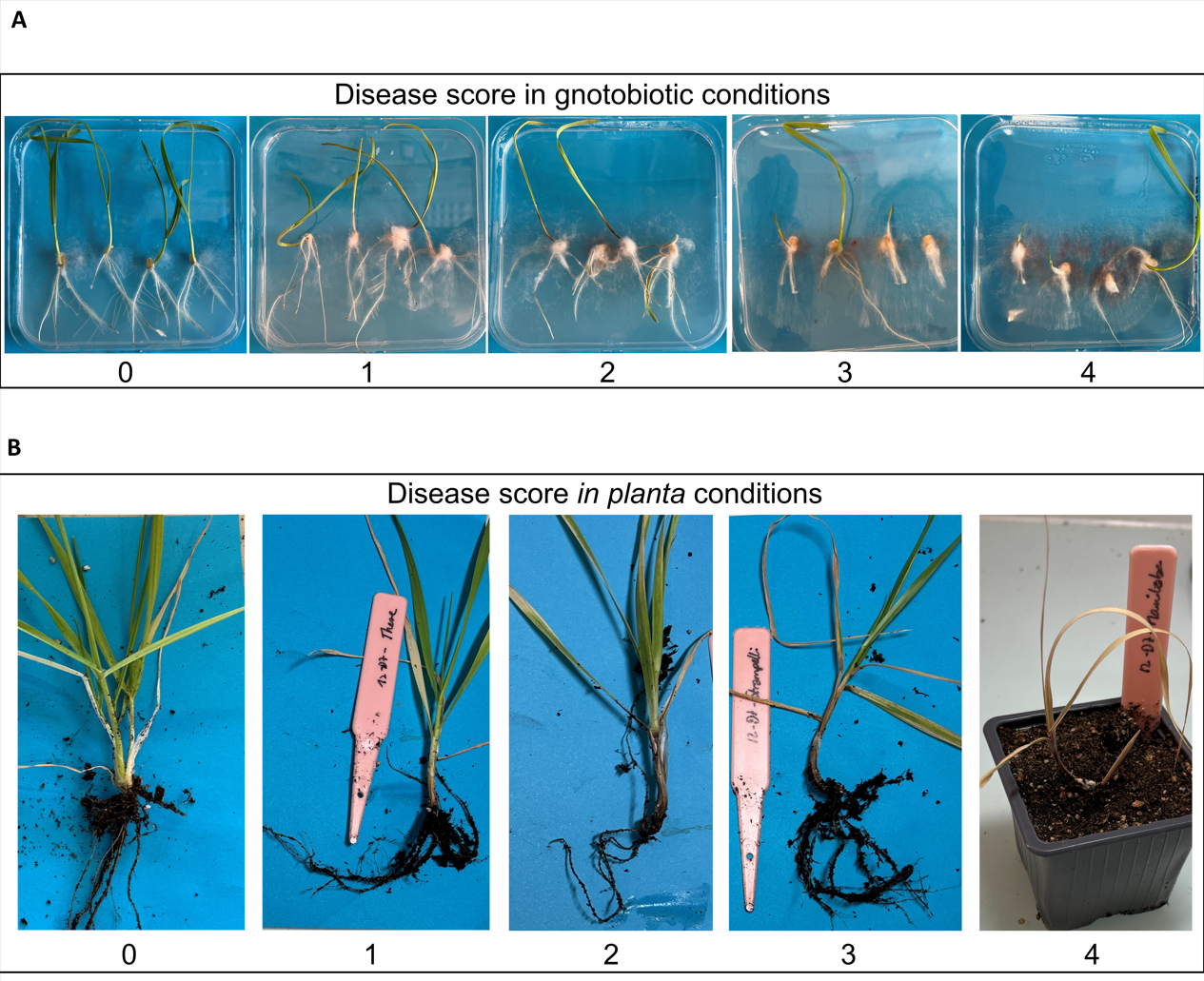

**Figure S1. Disease scale used in gnotobiotic and *in planta* conditions to evaluate the aggressiveness of *Fusarium* strains**. Plants were recorded one week post inoculation in gnotobiotic conditions (A), while disease records were measured 4 weeks after inoculation under *in planta* conditions (B).

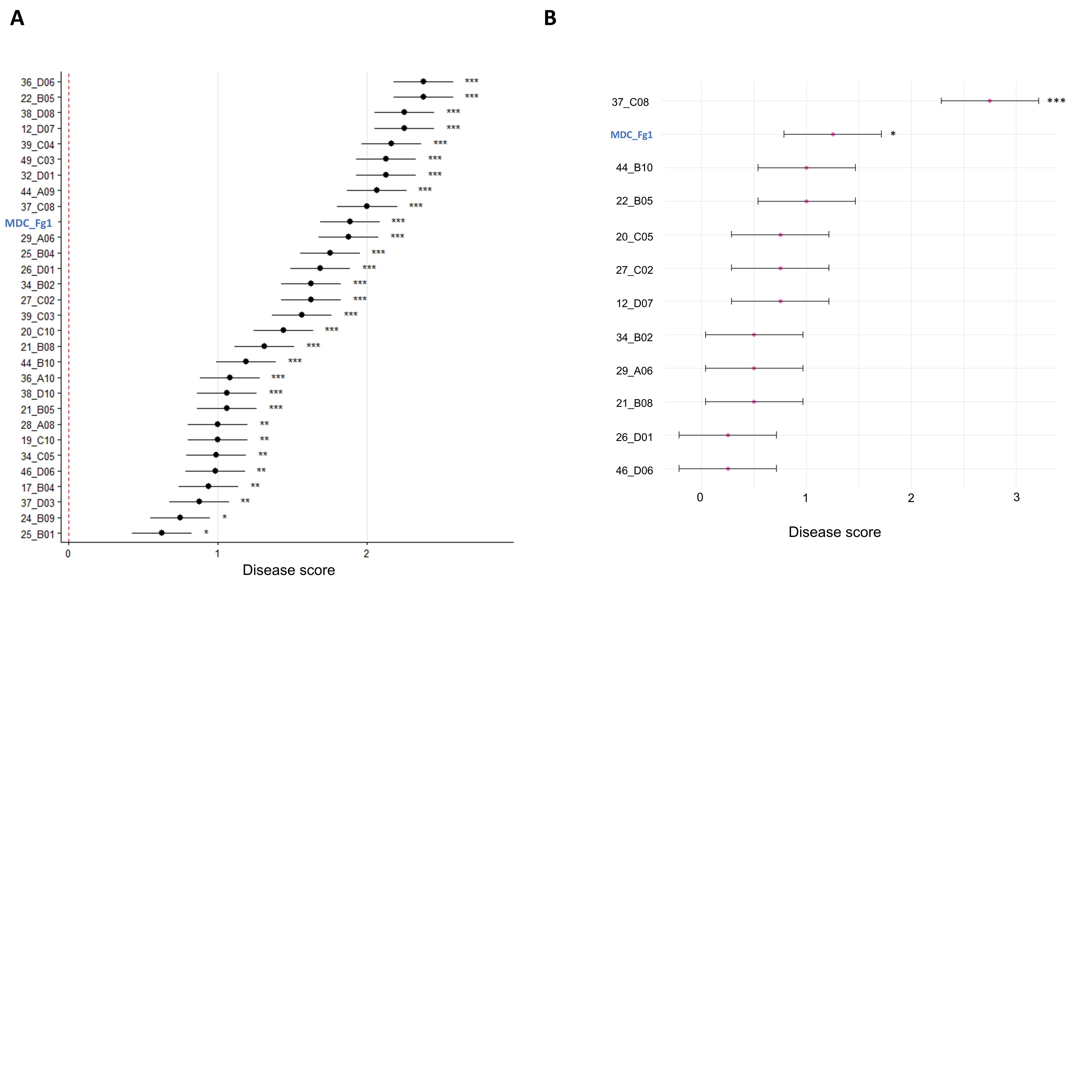

**Figure S2. Pathogenicity tests of *Fusarium* strains inoculated on the Paledor wheat variety.** (A) LS-means estimated using the disease score values of the 30 *Fusarium* strains causing significant Fusarium crown rot (FCR) symptoms under gnotobiotic conditions. The reference MDC_Fg1 is indicated in bold blue. The vertical red line indicates the negative control, corresponding to plants not inoculated with *Fusarium*. (B) LS-means for the *in planta* conditions pathogenicity test. For both tests only strains showing visible FCR symptoms are showed. Statistical significance is indicated as follows: *** *P* ≤ 0.0005, ** *P* ≤ 0.005, and * *P* ≤ 0.05.

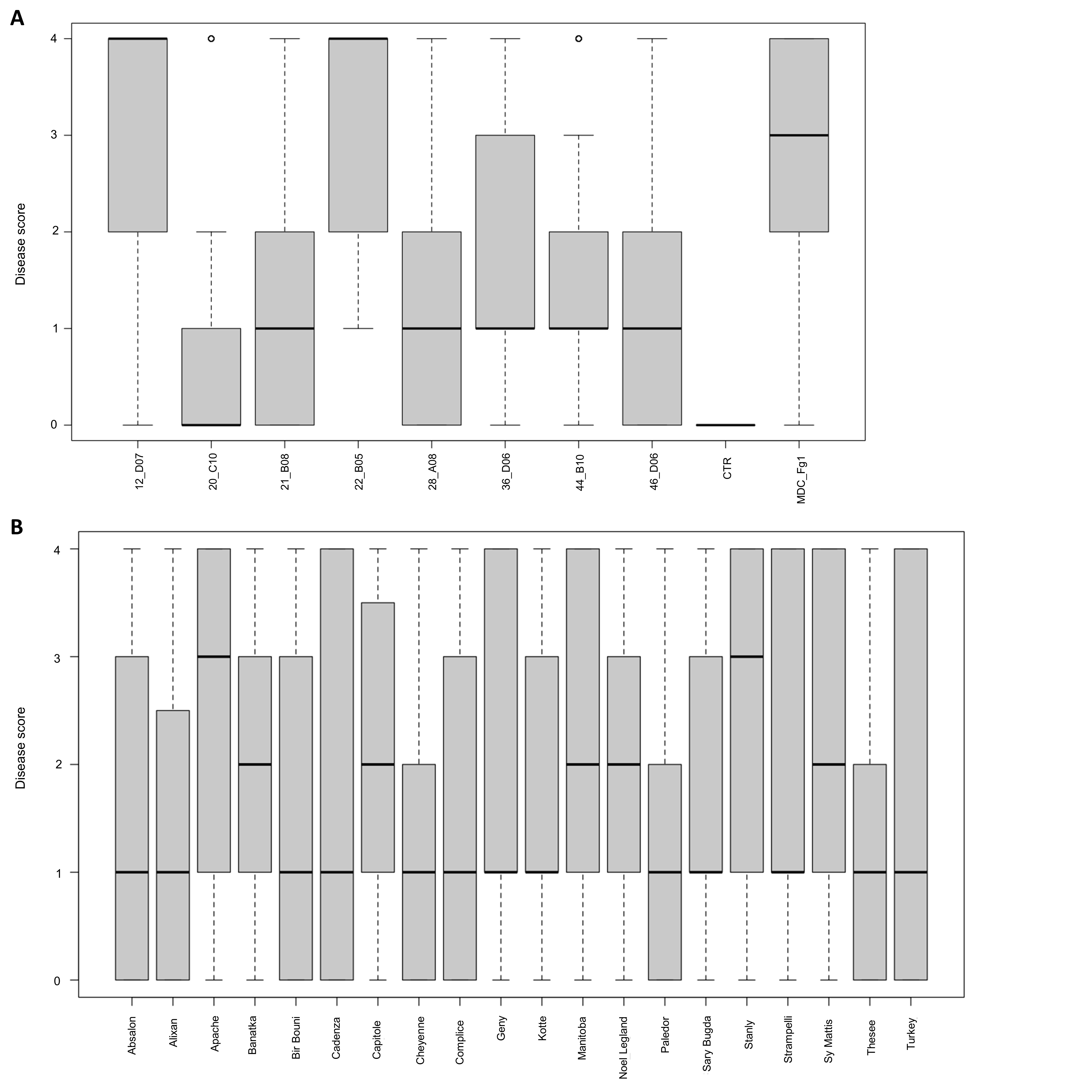

**Figure S3. Pathogenicity tests of eight *Fusarium* strains from non-agricultural habitats and the reference MDC_Fg1 on 20 wheat varieties.** (A) Box plot of the level of FCR disease (*y*-axis) distributed over the nine strains tested and the non-inoculated control (CTR) (*x*-axis). (B) Box plot of the level of FCR disease (*y*-axis) distributed over the 20 wheat varieties (*x*-axis).

**Figure S4. Plots of the LS-means estimated on the disease score values of eight *Fusarium* strains from non-agricultural habitats and the reference MDC_Fg1 on 20 wheat varieties.** (A) LS-means plot of the level of FCR disease (*y*-axis) distributed over the nine strains tested and the non-inoculated control (CTR) (*x*-axis). (B) LS-means plot of the level of FCR disease (*y*-axis) distributed over the 20 wheat varieties.

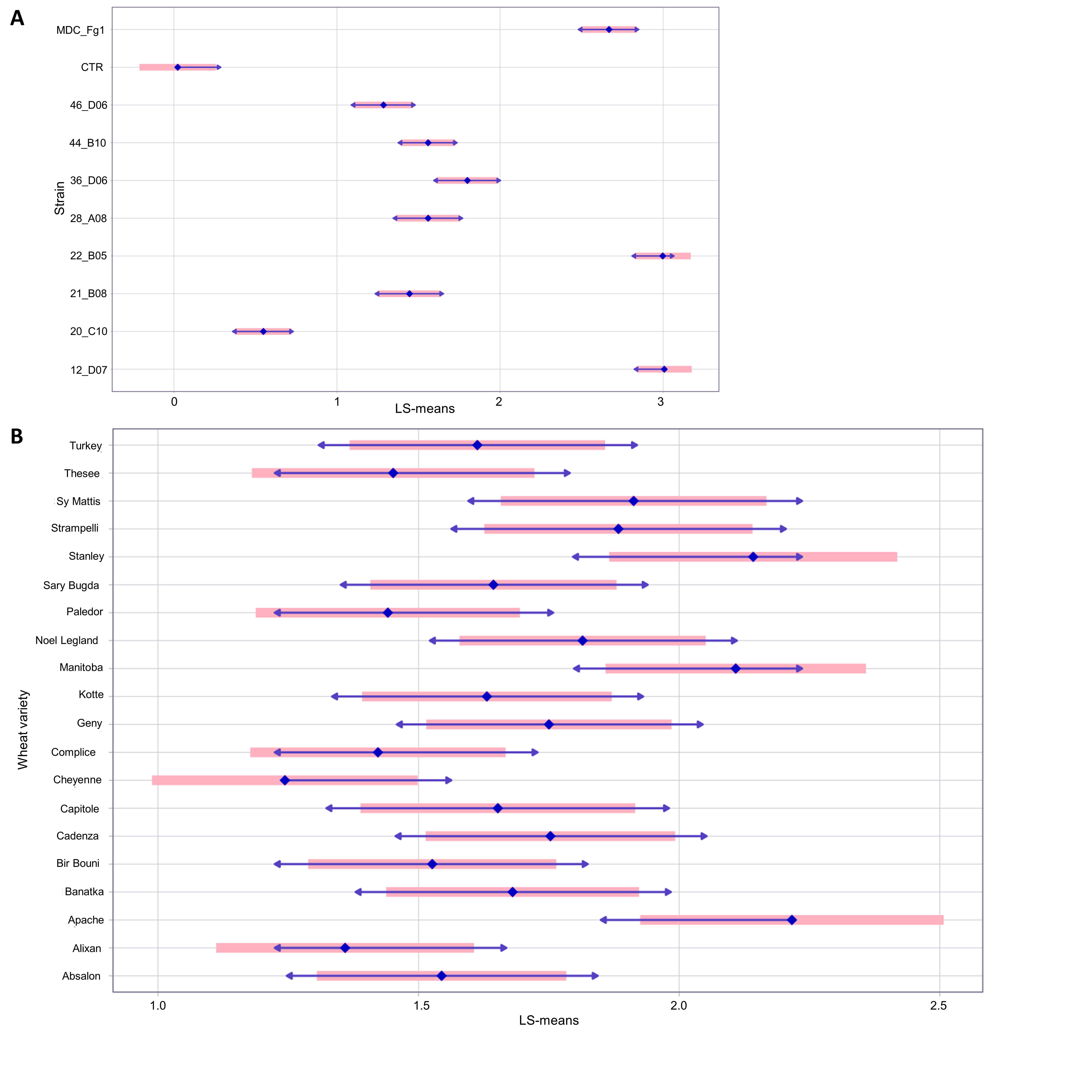

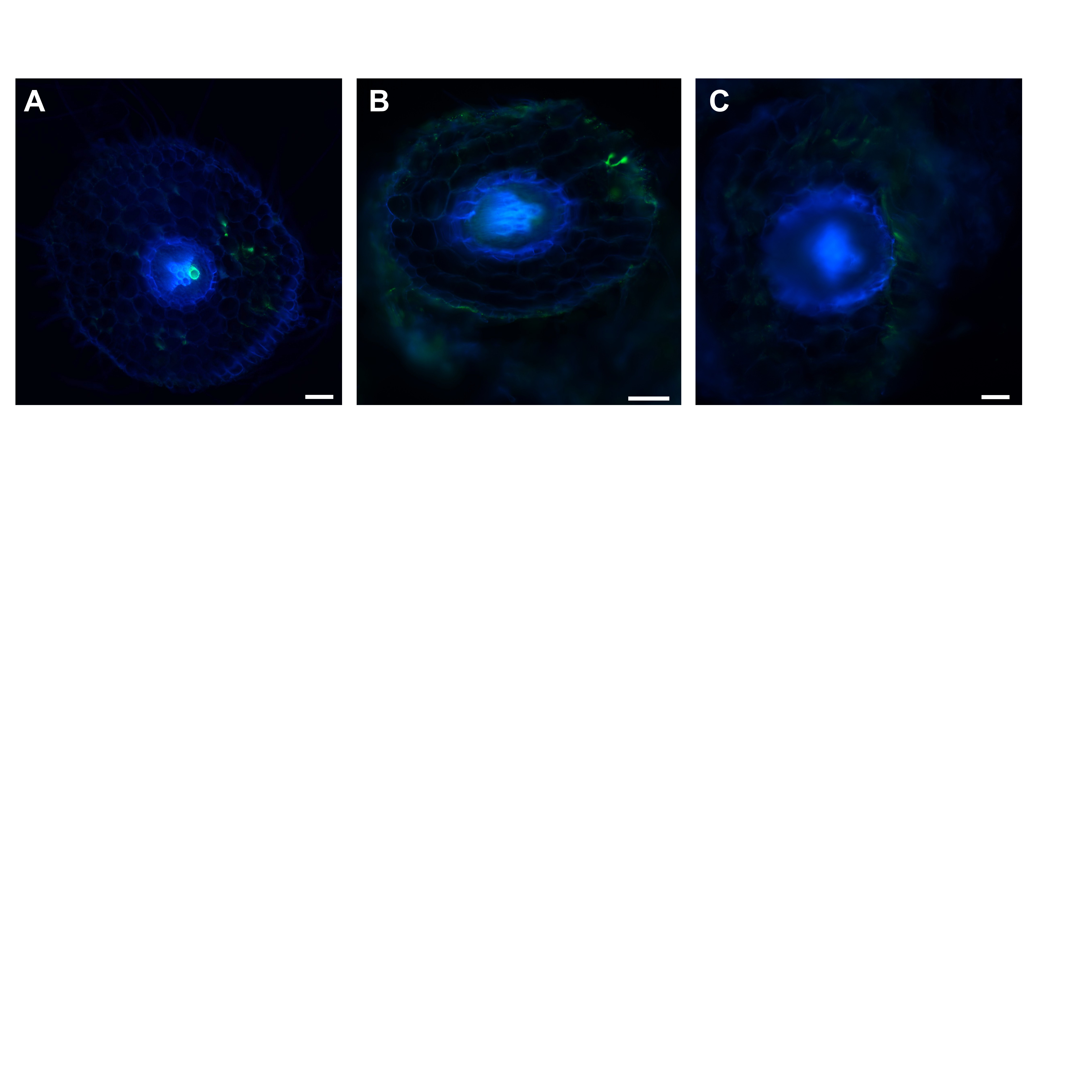

**Figure S5. Microscopy observation of *Fusarium* non-agricultural lineages in the *B. rapa* origin host.** Images of the colonization by the *Fusarium* strain 44_B10 (A-B) at 1-week post-inoculation (wpi) on *B. rapa*. After fungal staining with WGA-FITC, root sections were performed using a microtome and imaged by using a fluorescence microscope with a DAPI (blue, plant autofluorescence) and a GFP filter (green, fungal staining). Non-inoculated plants (C) were used as controls. Bars represent 50 µm.

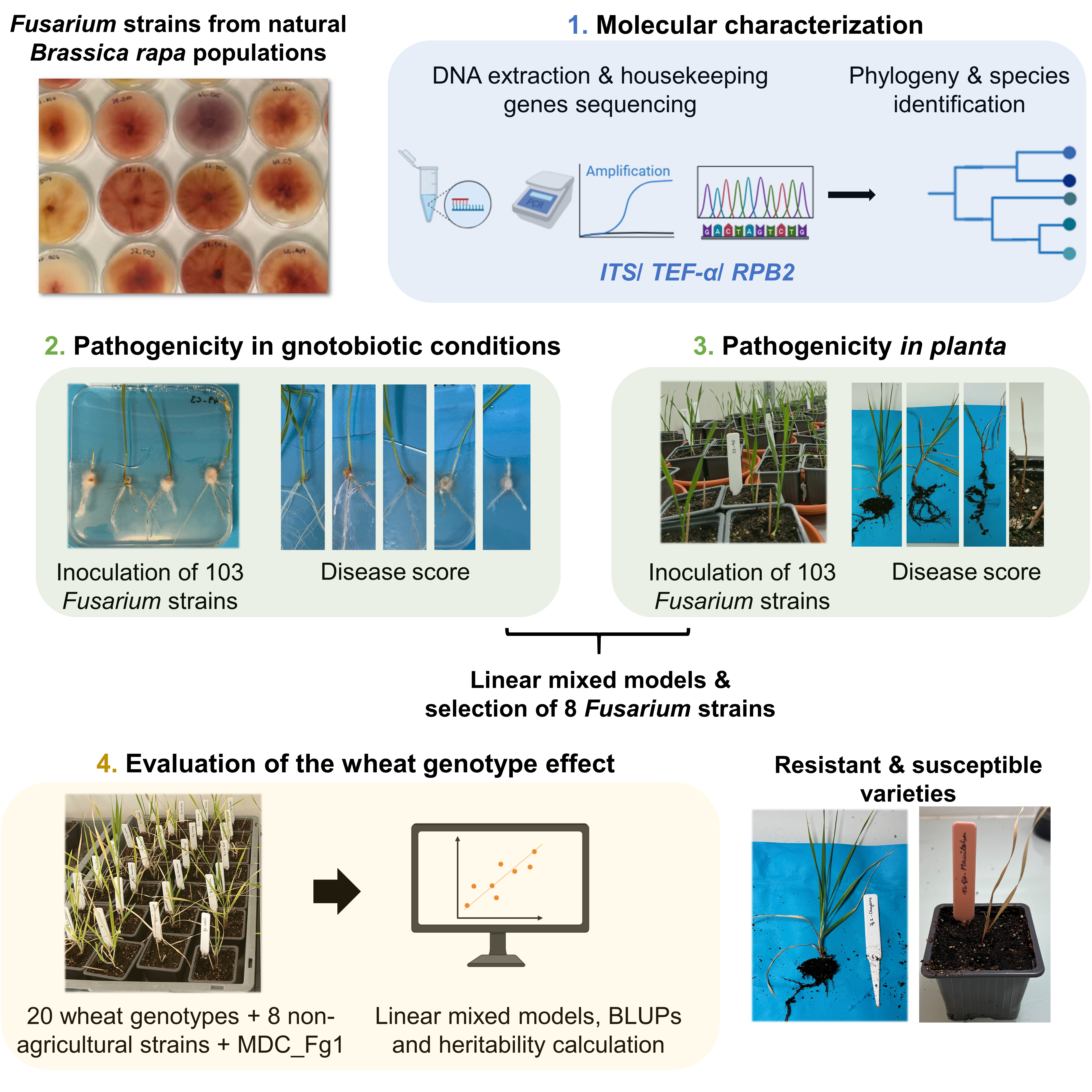

**Figure S6. Illustration of the experimental design adopted in our work**. *Fusarium* isolated strains were phylogenetically characterized and their pathogenicity was evaluated in gnotobiotic and planta conditions. The 103 *Fusarium* strains referred to the 102 non-agricultural strains and the reference MDC_Fg1. A set of 8 non-agricultural strains and the reference *F. graminearum* MDC_Fg1 strain were evaluated for their pathogenicity on a wheat core collection of 20 varieties.

**Supplementary Tables**

**Tables S1**. List of *Fusarium* strains isolated from *Brassica rapa* natural populations used in this study**.**

| **Strain code** | **Strain name** | **Country** | **City** | **GPS coordinates** |
| --- | --- | --- | --- | --- |
| 03_A08 | MARI_A_SP_RH02_2F_09 | Italy | Marineo | 37°58'07.9920"N 13°26'31.5780"E |
| 03_A10 | MARI_A_SP_RH01_2F_07 | Italy | Marineo | 37°58'07.9920"N 13°26'31.5780"E |
| 03_D06 | MARI_A_SP_RH02_2F_04 | Italy | Marineo | 37°58'07.9920"N 13°26'31.5780"E |
| 03_D07 | MARI_A_SP_RH02_2F_01 | Italy | Marineo | 37°58'07.9920"N 13°26'31.5780"E |
| 04_C02 | GRAT_A_SP_RH01_3F_04 | Italy | Gratteri | 37°58'04.1"N 13°58'52.6"E |
| 04_C03 | GRAT_A_SP_RH02_3F_01 | Italy | Gratteri | 37°58'04.1"N 13°58'52.6"E |
| 05_B02 | BOLO_A_SP_RH02_4F_02 | Italy | Bolognetta | 37°58'59.2920"N 13°26'26.0760"E |
| 05_B04 | BOLO_A_SP_RH01_4F_04 | Italy | Bolognetta | 37°58'59.2920"N 13°26'26.0760"E |
| 06_C05 | MORD_A_AT_RH08_4F_02 | France | Mordelles | 48°04'17.2"N 1°52'34.9"W |
| 07_D07 | GRAT_B_SP_RH10_6F_02 | Italy | Gratteri | 37°58'41.2"N 13°58'35.7"E |
| 08_A06 | MARI_A_SP_RH02_6F_03 | Italy | Marineo | 37°58'07.9920"N 13°26'31.5780"E |
| 08_C09 | MARI_A_SP_RH02_7F_07 | Italy | Marineo | 37°58'07.9920"N 13°26'31.5780"E |
| 08_D10 | ISNE_B_SP_RH01_7F_01 | Italy | Galefina | 37°56'34.4"N 14°01'01.1"E |
| 09_C08 | MARI_A_SP_RH02_7F_03 | Italy | Marineo | 37°58'07.9920"N 13°26'31.5780"E |
| 09_C09 | MARI_A_SP_RH02_7F_04 | Italy | Marineo | 37°58'07.9920"N 13°26'31.5780"E |
| 12_D07 | MORD_A_AT_RH10_9F_02 | France | Mordelles | 48°04'17.2"N 1°52'34.9"W |
| 13_C03 | MARI_A_SP_RH02_10F_03 | Italy | Marineo | 37°58'07.9920"N 13°26'31.5780"E |
| 17_B04 | MORD_A_AT_RH09_15F_03 | France | Mordelles | 48°04'17.2"N 1°52'34.9"W |
| 17_D06 | MARI_A_SP_RH02_15F_09 | Italy | Marineo | 37°58'07.9920"N 13°26'31.5780"E |
| 19_C10 | MORD_A_AT_RH10_1F_52 | France | Mordelles | 48°04'17.2"N 1°52'34.9"W |
| 20_B05 | MORD_A_AT_RH08_2F_01 | France | Mordelles | 48°04'17.2"N 1°52'34.9"W |
| 20_B10 | FOUG_A_AT_RH08_2F_01 | France | Fougeré | 47°37'59.0"N 0°08'51.7"W |
| 20_C05 | MORD_A_AT_R09_3F_02 | France | Mordelles | 48°04'17.2"N 1°52'34.9"W |
| 20_C09 | MORD_A_AT_R10_3F_01 | France | Mordelles | 48°04'17.2"N 1°52'34.9"W |
| 20_C10 | MORD_A_AT_RH10_3F_02 | France | Mordelles | 48°04'17.2"N 1°52'34.9"W |
| 20_D05 | ISNE_A_SP_RH01_3F_02 | Italy | Vinziria | 37°56'31.3"N 14°03'48.0"E |
| 21_A06 | FOUG_A_AT_RH09_2F_01 | France | Fougeré | 47°37'59.0"N 0°08'51.7"W |
| 21_B05 | RENN_A_AT_R08_1F_02 | France | Rennes | 48°06'22.9"N 1°42'33.4"W |
| 21_B08 | MORD_A_AT_RH08_2F_02 | France | Mordelles | 48°04'17.2"N 1°52'34.9"W |
| 21_D03 | POLL_A_SP_RH01_2F_02 | Italy | Pollina | 37°57' 57.24''N 14° 6' 9.705''E |
| 22_A03 | MORD_A_AT_L10_3F_01 | France | Mordelles | 48°04'17.2"N 1°52'34.9"W |
| 22_B05 | BRUZ_A_AT_RH10_3F_01 | France | Bruz | 48°02'30.1"N 1°46'34.0"W |
| 22_D06 | MORD_A_AT_R10_4F_04 | France | Mordelles | 48°04'17.2"N 1°52'34.9"W |
| 24_A03 | BOEL_A_AT_L10_4F_02 | France | Boël | 47°58’57.5"N  1°45’20.4"W |
| 24_B09 | MORD_A_AT_L10_4F_02 | France | Mordelles | 48°04'17.2"N 1°52'34.9"W |
| 24_C05 | BISA_A_SP_RH01_13F_01 | Italy | Bisacquino | 37°41'27.7560"N 13°16'47.6700"E |
| 25_A08 | CAST_A_SP_RH01_5F_01 | Italy | Castelbuono (Pollina) | 37°58' 38.964''N 14° 06' 8.712''E |
| 25_B01 | MORD_A_AT_RH10_5F_03 | France | Mordelles | 48°04'17.2"N 1°52'34.9"W |
| 25_B04 | APIG_A_AT_RH08_5F_03 | France | Etangs d'Apigné | 48°05'37.0"N 1°44'47.8"W |
| 26_D01 | BRUZ_A_AT_RH08_6F_01 | France | Bruz | 48°02'30.1"N 1°46'34.0"W |
| 26_D04 | MORD_A_AT_R09_6F_02 | France | Mordelles | 48°04'17.2"N 1°52'34.9"W |
| 27_B07 | POLL_A_SP_RH02_7F_03 | Italy | Pollina | 37°57' 57.24''N 14° 6' 9.705''E |
| 27_C02 | BRUZ_A_AT_R08_7F_04 | France | Bruz | 48°02'30.1"N 1°46'34.0"W |
| 28_A08 | MORD_A_AT_RH10_7F_53 | France | Mordelles | 48°04'17.2"N 1°52'34.9"W |
| 28_B03 | POLL_A_SP_RH02_7F_01 | Italy | Pollina | 37°57' 57.24''N 14° 6' 9.705''E |
| 28_B05 | FOUG_A_AT_R10_7F_01 | France | Fougeré | 47°37'59.0"N 0°08'51.7"W |
| 29_A06 | BRUZ_A_AT_RH10_8F_02 | France | Bruz | 48°02'30.1"N 1°46'34.0"W |
| 29_C05 | MISI_B_SP_RH01_8F_03 | Italy | Misilmeri | 38°00'09.8040"N 13°26'44.7060"E |
| 30_A01 | CAST_B_SP_RH01_9F_02 | Italy | Castelbuono | 37°56'53.3"N 14°05'38.2"E |
| 30_B05 | POLL_A_SP_RH02_9F_02 | Italy | Pollina | 37°57' 57.24''N 14° 6' 9.705''E |
| 32_B01 | CAST_B_SP_RH02_14F_01 | Italy | Castelbuono | 37°56'53.3"N 14°05'38.2"E |
| 32_C01 | MORD_A_AT_R09_14F_04 | France | Mordelles | 48°04'17.2"N 1°52'34.9"W |
| 32_C06 | BRUZ_A_AT_RH10_14F_05 | France | Bruz | 48°02'30.1"N 1°46'34.0"W |
| 32_D01 | BOEL_A_AT_RH08_14F_03 | France | Boël | 47°58’57.5"N  1°45’20.4"W |
| 33_A05 | ISNE_A_SP_RH01_14F_02 | Italy | Vinziria | 37°56'31.3"N 14°03'48.0"E |
| 33_A07 | RENN_A_AT_R08_14F_02 | France | Rennes | 48°06'22.9"N 1°42'33.4"W |
| 33_D05 | POLL_A_SP_RH01_15F_02 | Italy | Pollina | 37°57' 57.24''N 14° 6' 9.705''E |
| 34_B02 | BRUZ_A_AT_RH10_15F_03 | France | Bruz | 48°02'30.1"N 1°46'34.0"W |
| 34_B08 | MISI_A_SP_RH01_16F_01 | Italy | Misilmeri | 38°01'04.0860"N 13°26'52.5960"E |
| 34_C05 | BRUZ_A_AT_RH10_16F_02 | France | Bruz | 48°02'30.1"N 1°46'34.0"W |
| 35_C02 | BRUZ_A_SP_RH_2F_03 | France | Bruz | 48°02'30.1"N 1°46'34.0"W |
| 35_D01 | BRUZ_A_SP_RH_3F_02 | France | Bruz | 48°02'30.1"N 1°46'34.0"W |
| 36_A10 | BOEL_A_SP_RH_4F_01 | France | Boël | 47°58’57.5"N  1°45’20.4"W |
| 36_C07 | BRUZ_A_SP_RH_5F_01 | France | Bruz | 48°02'30.1"N 1°46'34.0"W |
| 36_C08 | BRUZ_A_SP_RH_5F_02 | France | Bruz | 48°02'30.1"N 1°46'34.0"W |
| 36_D06 | MORD_A_SP_RH_5F_04 | France | Mordelles | 48°04'17.2"N 1°52'34.9"W |
| 37_A05 | BRUZ_A_SP_RH_6F_01 | France | Bruz | 48°02'30.1"N 1°46'34.0"W |
| 37_C08 | FOUG_A_SP_R_9F_01 | France | Fougeré | 47°37'59.0"N 0°08'51.7"W |
| 37_D01 | BOEL_A_SP_RH_8F_01 | France | Boël | 47°58’57.5"N  1°45’20.4"W |
| 37_D03 | BOEL_A_SP_RH_8F_03 | France | Boël | 47°58’57.5"N  1°45’20.4"W |
| 37_D05 | BRUZ_A_SP_RH_8F_01 | France | Bruz | 48°02'30.1"N 1°46'34.0"W |
| 38_A07 | BOEL_A_SP_RH_10F_01 | France | Boël | 47°58’57.5"N  1°45’20.4"W |
| 38_B07 | BRUZ_A_SP_RH_10F_01 | France | Bruz | 48°02'30.1"N 1°46'34.0"W |
| 38_D04 | BOEL_A_SP_RH_14F_03 | France | Boël | 47°58’57.5"N  1°45’20.4"W |
| 38_D08 | RENN_A_SP_R_14F_02 | France | Rennes | 48°06'22.9"N 1°42'33.4"W |
| 38_D10 | BOEL_A_SP_RH_15F_03 | France | Boël | 47°58’57.5"N  1°45’20.4"W |
| 39_A04 | BRUZ_A_SP_RH_15F_02 | France | Bruz | 48°02'30.1"N 1°46'34.0"W |
| 39_C03 | BOEL_A_SP_RH_15F_02 | France | Boël | 47°58’57.5"N  1°45’20.4"W |
| 39_C04 | RENN_A_SP_RH_14F_03 | France | Rennes | 48°06'22.9"N 1°42'33.4"W |
| 40_B04 | PIAN_A_AT_RH_1F_03 | Italy | Piana degli Albanesi | 38°01'13"N 13°18'47"E |
| 40_C02 | BISA_A_AT_RH_1F_03 | Italy | Bisacquino | 37°41'27.7560"N 13°16'47.6700"E |
| 41_A01 | MARI_A_AT_RH_2F_03 | Italy | Marineo | 37°58'07.9920"N 13°26'31.5780"E |
| 41_A05 | LASC_A_AT_RH_2F_01 | Italy | Lascari | 37°59'12.5"N 13°56'26.6"E |
| 41_B02 | ALTO_A_AT_RH_2F_03 | Italy | Altofonte | 38°02'18"N 13°19'04"E |
| 41_C05 | BISA_A_AT_RH_3F_05 | Italy | Bisacquino | 37°41'27.7560"N 13°16'47.6700"E |
| 42_A07 | ISNE_A_AT_RH_4F_01 | Italy | Vinziria | 37°56'31.3"N 14°03'48.0"E |
| 43_A07 | CAST_A_AT_RH_4F_04 | Italy | Castelbuono (Pollina) | 37°58' 38.964''N 14° 06' 8.712''E |
| 44_A09 | CAST_A_AT_RH_6F_01 | Italy | Castelbuono (Pollina) | 37°58' 38.964''N 14° 06' 8.712''E |
| 44_B10 | GRAT_A_AT_RH_7F_01 | Italy | Gratteri | 37°58'04.1"N 13°58'52.6"E |
| 44_C05 | GRAT_B_AT_RH_7F_02 | Italy | Gratteri | 37°58'41.2"N 13°58'35.7"E |
| 45_D04 | BISA_A_AT_RH_10F_02 | Italy | Bisacquino | 37°41'27.7560"N 13°16'47.6700"E |
| 46_C02 | GRAT_B_AT_RH_12F_02 | Italy | Gratteri | 37°58'41.2"N 13°58'35.7"E |
| 46_D06 | COLL_A_AT_RH_13F_04 | Italy | Collesano | 37°51'01"N 13°54'49"E |
| 47_A07 | BISA_A_AT_RH_13F_02 | Italy | Bisacquino | 37°41'27.7560"N 13°16'47.6700"E |
| 47_C04 | CORL_A_AT_RH_14F_02 | Italy | Corleone | 37°48'03.7860"N 13°18'00.3900"E |
| 48_B04 | CAST_C_AT_RH_15F_04 | Italy | Castelbuono | 37°56'24.6"N 14°05'33.9"E |
| 48_B06 | MARI_A_AT_RH_16F_02 | Italy | Marineo | 37°58'07.9920"N 13°26'31.5780"E |
| 48_D04 | MONR_A_AT_RH_15F_03 | Italy | Monreale | 38°01'02"N 13°15'01"E |
| 49_B06 | MARI_A_AT_R_10F_05 | Italy | Marineo | 37°58'07.9920"N 13°26'31.5780"E |
| 49_C01 | FOUG_A_SP_RH_14F_02 | France | Fougeré | 47°37'59.0"N 0°08'51.7"W |
| 49_C03 | FOUG_A_SP_RH_16F_02 | France | Fougeré | 47°37'59.0"N 0°08'51.7"W |
| 49_C07 | MARI_A_AT_RH_7F_05 | Italy | Marineo | 37°58'07.9920"N 13°26'31.5780"E |

**Table S2**. List of 20 wheat varieties used to test the effect of host genotype of Fusarium crown rot (FCR) disease of eight non-agricultural *Fusarium* strains and the reference *F. graminearum* MDC_Fg1. NA: not available.

| **Name of the variety** | **Type** | **Date of inscription** |
| --- | --- | --- |
| Absalon | Modern | 2016 |
| Alixan | Modern | 2005 |
| Apache | Modern | 1998 |
| Banatka | Ancient | NA |
| Bir Bouni | Ancient | NA |
| Cadenza | Modern | 1995 |
| Capitole | Modern | 1963 |
| Cheyenne | Ancient | 1933 |
| Complice | Modern | 2016 |
| Geny | Ancient | 2019 |
| Kotte | Ancient | 1896 |
| Manitoba | Ancient | 1923 |
| Noel-Legland | Ancient | 1940 |
| Paledor | Modern | 2005 |
| Sary-Bugda | Ancient | NA |
| Stanley | Ancient | 1890 |
| Strampelli | Ancient | 1927 |
| SY Matthis | Modern | 2010 |
| Thésée | Modern | 1983 |
| Turkey | Ancient | 1874 |

**Table S3**. Pairwise comparisons between the nine *Fusarium* strains and the negative control (CTR) on 20 wheat varieties five weeks post-inoculation.

| **Contrast** | **Estimate** | **SE** | **T ratio** | ***P* value** |
| --- | --- | --- | --- | --- |
| 12_D7 - CTR | 2.9827216 | 0.1415615 | 21.070 | <0.0001 |
| 20_C10 - CTR | 0.5250204 | 0.1427430 | 3.678 | 0.0002 |
| 21_B08 - CTR | 1.4208449 | 0.1490897 | 9.530 | <0.0001 |
| 22_B5 - CTR | 2.9732533 | 0.1427442 | 20.829 | <0.0001 |
| 28_A08 - CTR | 1.5351230 | 0.1505142 | 10.199 | <0.0001 |
| 36_D06 - CTR | 1.7757973 | 0.1476666 | 12.026 | <0.0001 |
| 44_B10 - CTR | 1.5351115 | 0.1414096 | 10.856 | <0.0001 |
| 46_D06 - CTR | 1.2615512 | 0.1455101 | 8.670 | <0.0001 |
| MDC_Fg1 - CTR | 2.6440437 | 0.1425916 | 18.543 | <0.0001 |

**Table S4**. Pairwise comparisons between the nine *Fusarium* strains and the reference MDC_Fg1 used as a positive control on 20 wheat five weeks post-inoculation.

| **Contrast** | **Estimate** | **SE** | **T ratio** | ***P* value** |
| --- | --- | --- | --- | --- |
| 12_D7 - MDC_Fg1 | 0.3386778 | 0.1155520 | 2.931 | 0.0038 |
| 20_C10 - MDC_Fg1 | -2.1190233 | 0.1170215 | -18.108 | <0.0001 |
| 21_B08 - MDC_Fg1 | -1.2231988 | 0.1250465 | -9.782 | <0.0001 |
| 22_B5 - MDC_Fg1 | 0.3292096 | 0.1169360 | 2.815 | 0.0049 |
| 28_A08 - MDC_Fg1 | -1.1089207 | 0.1260102 | -8.800 | <0.0001 |
| 36_D06 - MDC_Fg1 | -0.8682464 | 0.1234062 | -7.036 | <0.0001 |
| 44_B10 - MDC_Fg1 | -1.1089322 | 0.1152805 | -9.619 | <0.0001 |
| 46_D06 - MDC_Fg1 | -1.3824925 | 0.1205451 | -11.469 | <0.0001 |
| CTR - MDC_Fg1 | -2.6440437 | 0.1425916 | -18.543 | <0.0001 |

**Table S5**. Pairwise comparisons between *Fusarium* strains during the experimental assays testing the development of FCR disease in the 20 wheat varieties, at five weeks post-inoculation.

| **Contrast** | **Estimate** | **SE** | **T ratio** | ***P* value** |
| --- | --- | --- | --- | --- |
| 12_D7-20_C10 | 2.46e+00 | 0.116 | 21.251 | <0.0001 |
| 12_D7-21_B08 | 1.56e+00 | 0.124 | 12.616 | <0.0001 |
| 12_D7-22_B5 | 9.47e-03 | 0.116 | 0.082 | 1.0000 |
| 12_D7-28_A08 | 1.45e+00 | 0.125 | 11.603 | <0.0001 |
| 12_D7-36_D06 | 1.21e+00 | 0.122 | 9.879 | <0.0001 |
| 12_D7-44_B10 | 1.45e+00 | 0.114 | 12.710 | <0.0001 |
| 12_D7-46_D06 | 1.72e+00 | 0.119 | 14.432 | <0.0001 |
| 12_D7-CTR | 2.98e+00 | 0.142 | 21.070 | <0.0001 |
| 12_D7-MDC_Fg1 | 3.39e-01 | 0.116 | 2.931 | 0.0978 |
| 20_C10-21_B08 | -8.96e-01 | 0.125 | -7.162 | <0.0001 |
| 20_C10-22_B5 | -2.45e+00 | 0.117 | -20.901 | <0.0001 |
| 20_C10-28_A08 | -1.01e+00 | 0.126 | -8.010 | <0.0001 |
| 20_C10-36_D06 | -1.25e+00 | 0.123 | -10.131 | <0.0001 |
| 20_C10-44_B10 | -1.01e+00 | 0.115 | -8.754 | <0.0001 |
| 20_C10-46_D06 | -7.37e-01 | 0.121 | -6.109 | <0.0001 |
| 20_C10-CTR | 5.25e-01 | 0.143 | 3.678 | 0.0091 |
| 20_C10-MDC_Fg1 | -2.12e+00 | 0.117 | -18.108 | <0.0001 |
| 21_B08-22_B5 | -1.55e+00 | 0.125 | -12.397 | <0.0001 |
| 21_B08-28_A08 | -1.14e-01 | 0.133 | -0.859 | 0.9975 |
| 21_B08-36_D06 | -3.55e-01 | 0.131 | -2.719 | 0.1668 |
| 21_B08-44_B10 | -1.14e-01 | 0.124 | -0.925 | 0.9957 |
| 21_B08-46_D06 | 1.59e-01 | 0.128 | 1.245 | 0.9650 |
| 21_B08-CTR | 1.42e+00 | 0.149 | 9.530 | <0.0001 |
| 21_B08-MDC_Fg1 | -1.22e+00 | 0.125 | -9.782 | <0.0001 |
| 22_B5-28_A08 | 1.44e+00 | 0.126 | 11.406 | <0.0001 |
| 22_B5-36_D06 | 1.20e+00 | 0.124 | 9.695 | <0.0001 |
| 22_B5-44_B10 | 1.44e+00 | 0.115 | 12.465 | <0.0001 |
| 22_B5-46_D06 | 1.71e+00 | 0.121 | 14.181 | <0.0001 |
| 22_B5-CTR | 2.97e+00 | 0.143 | 20.829 | <0.0001 |
| 22_B5-MDC_Fg1 | 3.29e-01 | 0.117 | 2.815 | 0.1319 |
| 28_A08-36_D06 | -2.41e-01 | 0.132 | -1.828 | 0.7171 |
| 28_A08-44_B10 | 1.14e-05 | 0.125 | 0.000 | 1.0000 |
| 28_A08-46_D06 | 2.74e-01 | 0.129 | 2.120 | 0.5135 |
| 28_A08-CTR | 1.54e+00 | 0.151 | 10.199 | <0.0001 |
| 28_A08-MDC_Fg1 | -1.11e+00 | 0.126 | -8.800 | <0.0001 |
| 36_D06-44_B10 | 2.41e-01 | 0.122 | 1.973 | 0.6180 |
| 36_D06-46_D06 | 5.14e-01 | 0.126 | 4.066 | 0.0020 |
| 36_D06-CTR | 1.78e+00 | 0.148 | 12.026 | <0.0001 |
| 36_D06-MDC_Fg1 | -8.68e-01 | 0.123 | -7.036 | <0.0001 |
| 44_B10-46_D06 | 2.74e-01 | 0.119 | 2.298 | 0.3905 |
| 44_B10-CTR | 1.54e+00 | 0.141 | 10.856 | <0.0001 |
| 44_B10-MDC_Fg1 | -1.11e+00 | 0.115 | -9.619 | <0.0001 |
| 46_D06-CTR | 1.26e+00 | 0.146 | 8.670 | <0.0001 |
| 46_D06-MDC_Fg1 | -1.38e+00 | 0.121 | -11.469 | <0.0001 |
| CTR-MDC_Fg1 | -2.64e+00 | 0.143 | -18.543 | <0.0001 |

**Table S6**. Primers used to amplify housekeeping genes in *Fusarium* strains.

| **Locus** | **Primer name** | **Sequence 5’-3’** |
| --- | --- | --- |
| ***TEF-α*** | EF1 | ATGGGTAAGGARGACAAGAC |
|  | EF2 | GGARGTACCAGTSATCATGTT |
| ***RPB2*** | 5f2 | GGGGWGAYCAGAAGAAGGC |
|  | 7cr | CCCATRGCTTGYTTRCCCAT |
| ***ITS1*** | ITS1Fngs | GGTCATTTAGAGGAAGTAA |
|  | ITS2Fngs | TTYRCKRCGTTCTTCATCG |

**Supplementary Text**

**Supplementary experimental procedures**

**Molecular characterization of *Fusarium* strains isolates from non-agricultural habitats and phylogenetic analysis**

To determine the phylogenetic and taxonomic affiliation of the 102 *Fusarium* strains, DNA was extracted from each strain (Figure S6). For this purpose, strains were first cultured on PDA plates. After one week of growth, the mycelium was scraped from the surface of the plates and transferred to a 2-mL Eppendorf tube. The fungal mycelium was weighed to obtain between 0.05 and 0.1 g of fresh mycelium. After adding a 0.5-cm-diameter ceramic bead, each tube was immersed in liquid nitrogen, and the fungal material was subsequently ground at 1800 rpm using a FastPrep® homogenizer for three 30-s cycles. DNA extraction was performed according to the protocol developed by Mayjonnade et al. (in preparation). The purity and quality of the extracted DNA were assessed using a NanoDrop® spectrophotometer. DNA samples with concentrations above 100 ng/µL were diluted tenfold.

Two microlitres of diluted DNA were used to amplify the housekeeping genes *TEF-1α*, *ITS1*, and *RPB2* (Table 2) (O’Donnell et al. 1998, 2022; Ma et al. 2024) using the GoTaq® PCR kit (Promega) (Figure S6, Table S6). The housekeeping genes were amplified by PCR using the following cycling conditions: an initial denaturation at 95°C for 5 min, followed by 36 cycles of 95°C for 30 s, 56°C for 1 min, and 72°C for 30 s, with a final extension step at 72°C for 10 min. Following electrophoresis of 5 µL of the PCR products on a 1% agarose gel for 30 min at 130 V, the gel was visualized using GelRed® to verify successful PCR amplification before Sanger sequencing by the Eurofins sequencing platform. The resulting sequences were subsequently analyzed using Chromas 2.6.6 (<https://technelysium.com.au/wp/chromas/>). The cleaned sequences were exported in FASTA format and analyzed using BLAST searches against the NCBI database to determine their potential taxonomic affiliation.

The phylogenetic analysis included the 102 *Fusarium* strains and the reference strain MDC_Fg1 was based on the three sequenced housekeeping genes. Reference sequences representing different *Fusarium* species were also retrieved from the NCBI and IMG-JGI databases. Reference sequences were selected based on the taxonomic identifications obtained from BLAST searches against GenBank. Some reference sequences could not be retrieved because sequence information for all three housekeeping genes was not available for certain species. The sequences were subsequently aligned using DAMBE (Xia 2018) with the MUSCLE alignment algorithm. The resulting alignments were used to construct a phylogenetic tree in MEGA 12 (Kumar et al. 2024), using the Neighbor-Joining method with a gamma distribution and 1,000 bootstrap replicates. Phenotypic data and phylogenetic relationships were subsequently visualized and analyzed using iTOL v7.6 (Letunic and Bork 2024).

**Statistical analyses to identify *Fusarium* strain aggressiveness and the effect of wheat genotype**

All statistical analyses and figures were generated using the R environment (version 4.5.3). To assess the pathogenicity of *Fusarium* strains under both gnotobiotic and *in planta* conditions, the following model was used:

*Disease* ~ *Strain* + (1 | *Rep*) + (1 | *Block*)

In the model implemented in the *lmer* R package, *Disease* corresponds to the disease index recorded for each plant, *Strain* corresponds to the *Fusarium* strain used (fixed factor), *Block* corresponds to the temporal replicate (random factor), and *Rep* corresponds to the plants within each temporal block (random factor). Adjusted means were subsequently estimated using the emmeans package, and *P* values were corrected using the False Discovery Rate (FDR) procedure. Only strains with an FDR-adjusted *P* value < 0.05 were considered pathogenic under gnotobiotic or *in planta* conditions.

To assess the effects of wheat variety and fungal strain on FCR disease severity, the following linear mixed-effects model was fitted using lmer:

*Disease* ~ *Variety* + *Strain* + (1 | *Rep*)

In this model, *Disease* corresponds to the disease index recorded for each plant, *Variety* corresponds to the wheat genotype (fixed factor), *Strain* corresponds to the *Fusarium* strain used (fixed factor), and *Rep* corresponds to the plants within each spatial block (random factor). Based on the significance of the effects in the overall model, pairwise comparisons were performed using adjusted means estimated with the emmeans package. Differences in pathogenicity among *Fusarium* strains and differences in disease resistance among wheat varieties were assessed using pairwise comparisons. *P* values were corrected using FDR. To further assess the contribution of wheat genotype to disease variation, broad-sense heritability was estimated using the inti package version 0.6.9, based on the following linear mixed-effects model:

*Disease* ~ *Strain* + (1 | *Variety*) + (1 | *Rep*)

In this model, *Disease* and *Strain* were treated as fixed factors, whereas *Variety* and *Rep* were treated as random factors. Given our experimental design, which included replicates and random spatial blocks, heritability was estimated from genotype BLUPs using the following equation:

𝐻^2^_𝐶𝑢𝑙𝑙𝑖𝑠_ = 1- $\frac{v\Delta- BLUP}{2*\sigma2g}$

where 𝑣∆-𝐵𝐿𝑈𝑃 is the mean squared error of the pairwise differences between genotypic BLUPs and 𝜎g is the genotypic variance.

**Bioinformatic analysis for the identification and characterization of *Fusarium*-affiliated OTUs obtained from *B. rapa* roots and rhizospheres**

The 102 *Fusarium* strains investigated in this study were isolated from the roots and rhizospheres of wild *B. rapa* populations collected in France (Brittany) and Italy (Sicily). Two complementary approaches were used to characterize the fungal communities associated with these samples. The first was a culture-dependent approach, which enabled the isolation of different fungal species, including members of the genus *Fusarium*. The second was a metabarcoding approach (Maillet et al. 2025). For the metabarcoding approach, environmental DNA extracted from roots and rhizospheres was amplified using the ITS marker to characterize the fungal community and obtain a description of the corresponding Operational Taxonomic Units (OTUs). In the present study, the metabarcoding data generated by Maillet et al. (2025) were retrieved and compared with the ITS sequences obtained from the 102 *Fusarium* strains isolated by culture-dependent methods. From these data, OTUs taxonomically affiliated with the genus *Fusarium* were extracted, and their relative abundance in each *B. rapa* population was calculated. This analysis identified six OTUs affiliated with *Fusarium*. The ITS sequences of these six OTUs were aligned with those of the 102 isolated *Fusarium* strains using DAMBE (Xia 2018). A phylogenetic tree was subsequently constructed using the Neighbor-Joining method to compare the metabarcoding-derived OTUs with the *Fusarium* strains isolated from *B. rapa* populations.

**Localization of *Fusarium* strains into wheat and *B. rapa* tissues**

To investigate the localization of non-agricultural *Fusarium* strains within plant tissues, we used the Manitoba spring wheat variety, the *B. rapa* BRUZ population (Maillet et al. 2025), and a fast-cycling *B. rapa* genotype used as a model for experimental evolution (Bartoli et al. 2026). Two non-agricultural *Fusarium* strains were tested on both host species: the highly aggressive strain 12_D07 and the moderately aggressive strain 44_B10. For each host and *B. rapa* genotype, *Fusarium* localization was assessed at four time points: 1-, 2-, and 3-weeks post-inoculation (wpi). At each time point, three plants were analysed for each host–strain combination, resulting in a total of six wheat plants and twelve *B. rapa* plants. Three plant compartments were analysed at each time point: roots, crown, and stem.

Before *Fusarium* inoculation, seeds of both plant species were surface-sterilized prior to germination, and *Fusarium* strains were inoculated as described for the pathogenicity assays. At each time point, the three plant compartments were collected, cut into sections, and immersed overnight in 100% ethanol at room temperature. The following day, the tissues were washed with PBS buffer, embedded in 4% agarose, and sectioned using a microtome at a thickness of 80 µm. The sections were placed in PBS until WGA-FITC (Invitrogen, Reference: W834) staining. For staining, PBS was removed and 1 mL of the stock solution (5mg/ml)1:1000-diluted WGA-FITC solution was added to the sections, which were incubated overnight at 4°C in the dark. The following day, the sections were washed three times with PBS before microscopic observation. Fluorescence and bright field images have been acquired on Nikon Eclipse TI inverted Wide field microscope with a colour CMOS camera DS Ri2 and drive by Nikon NIS software. In particular, we used 10x (N.A 0.3) or X20 ( N.A 0.45 ) objectives.
Fluorescence of WGA_FITC has been observed through a bandpass GFP B filter set (ex:475/50;dich:505;em:535/50) in order to visualize fungal hyphae, the autofluorescence of the cells walls by a bandpass DAPI filter set (ex:377/50;dich:409;em:447/60 ) to identify plant tissue structures.
